# An imbalance of naïve and effector T-cell phenotypes in early type 1 diabetes across conventional and regulatory subsets

**DOI:** 10.1101/2024.12.05.627068

**Authors:** Veronika Niederlova, Ales Neuwirth, Vit Neuman, Juraj Michalik, Bela Charvatova, Martin Modrak, Zdenek Sumnik, Ondrej Stepanek

**Affiliations:** Laboratory of Adaptive Immunity, Institute of Molecular Genetics of the Czech Academy of Sciences, Prague, Czechia; Department of Cell Biology, Faculty of Science, Charles University in Prague, Czechia; Department of Pediatrics, 2^nd^ Faculty of Medicine, Charles University in Prague & Motol University Hospital, Prague, Czechia; Department of Bioinformatics, Second Faculty of Medicine, Charles University, Prague, Czechia

**Keywords:** type 1 diabetes, self-tolerance, effector T cells, regulatory T cells, single-cell sequencing, Tr3-56 cells, hygiene hypothesis

## Abstract

Type 1 diabetes (T1D) is an autoimmune disease caused by the loss of self-tolerance toward insulin-producing pancreatic β-cells. Although the etiology of T1D is not fully understood, it is linked to dysregulation of the T-cell compartment. To identify T-cell signatures associated with T1D, we performed single-cell transcriptomic analysis of peripheral blood T-cells from newly diagnosed children, the same children after one year, and healthy donors. We observed reduced expression of genes related to effector and cytotoxic T-cell functions across conventional, unconventional, and regulatory T-cell subsets in diabetic children, particularly at the time of diagnosis. These findings were supported by flow cytometry analysis of the same cohort and by reanalysis of publicly available data. Overall, our results suggest that T1D is associated with impaired T-cell effector differentiation, which may contribute to immune dysregulation and loss of self-tolerance.

## Introduction

Type 1 diabetes mellitus (T1D) is an autoimmune disease characterized by the destruction of pancreatic β-cells, leading to insulin deficiency, hyperglycemia, and impaired metabolic homeostasis. As there is currently no cure for T1D, patients rely on insulin replacement therapy for the rest of their lives. Unlike most autoimmune diseases, T1D typically manifests in childhood. Both genetic and environmental factors contribute to the development of T1D. The genetic factors include specific HLA haplotypes ^1^, polymorphisms in the insulin gene (*INS*) ^2^ and several immune-related genes (e.g., *CTLA4*, *PTPN22*, *IFIH1*, and *CD226*) ^3^. The importance of the environmental factors is highlighted by a steep increase in T1D incidence in high-income countries ^4, 5, 6^. It has been proposed that specific viral infections, such as enteroviruses, can trigger T1D ^7, 8^. On the contrary, the so-called hygiene hypothesis explains the recent outburst of T1D and other autoimmune diseases by the reduction of the infection burden in children ^4, 9, 10, 11^. Despite some controversies in this area of research, it is well-established that a genetically and/or environmentally altered state of the immune system is a significant risk factor for T1D and may contribute directly to its onset. However, the etiology of T1D is still largely unknown.

T cells are extensively studied in the context of T1D because of the genetic evidence of HLA involvement, their clear role in the development of T1D in animal models ^12, 13^, and their presumed role in β-cell destruction via cell-mediated cytotoxicity ^14, 15^. Moreover, it has been proposed that T1D susceptibility may be induced by insufficient self-tolerance mediated by regulatory FOXP3^+^ T cells (Tregs) and non-classical regulatory T-cell populations, such as CD3^+^ CD56^+^ TR3-56 cells ^16^. Despite numerous studies in the field, the T1D-associated alterations in the T-cell compartment are still not fully understood ^17^.

Traditionally, the immune cells in T1D are studied using targeted flow cytometry panels or bulk transcriptomics on whole blood, peripheral blood mononuclear cells (PBMCs), and/or sorted leukocyte subsets ^18, 19, 20, 21, 22^. Whereas flow cytometry is limited to the pre-selected protein markers, bulk transcriptomic analysis lacks flow cytometry’s single cell resolution. These limitations can be overcome by multiparameter approaches on the single-cell level such as cytometry by time-of-flight or single cell RNA sequencing (scRNAseq). Indeed, pioneering studies established these techniques as the methods of choice to capture the differences between the immune cells in healthy donors and T1D children ^20, 23, 24, 25^. However, these studies also revealed potential caveats, such as a trade-off between the sufficient size of the cohort to address reproducibility and the number of sequenced cells per patient to achieve sufficient resolution ^20, 23, 25^. In parallel efforts, big consortia such as The Network for Pancreatic Organ Donors with Diabetes ^26^ and Human Pancreas Analysis Program (HPAP) ^27^ are generating large atlases of multimodal data from tissues of deceased donors, including scRNAseq, flow cytometry, immune repertoire profiling and imaging data. Although these efforts provide invaluable data sets including hardly accessible samples from solid organs, their disadvantage is their inability to recruit donors specifically based on the features such as their age or time after diagnosis. Moreover, a comprehensive analysis of scRNAseq data from the consortia focusing specifically on immune subsets has not been published yet.

To compare the T-cell compartments in healthy donors and T1D children and to monitor how T cells change in time in T1D children, we performed scRNAseq analysis of blood T cells of healthy donors and T1D patients at the time of diagnosis and again after one year. We used a balanced medium-sized cohort with a relatively high number of sequenced cells per donor. We observed a bias of conventional CD4^+^ and CD8^+^ T cells and unconvential CD8^+^ γδT cells towards a naïve or stem-like phenotype in newly diagnosed patients, which was partially normalized within one year after diagnosis. Moreover, the Treg-compartment in these patients showed a lack of functional differentiation and expressed a gene signature associated with dysfunction, in comparison to healthy donors. These conclusions were largely supported by our analyses of previously published data and by new analysis of publicly deposited data from independent cohorts.

## Results

### Generation of a T-cell atlas of children with T1D and healthy controls

We performed a prospective study on a medium-sized cohort of donors using scRNAseq to identify potential patterns in the gene expression and heterogeneity of blood T cells associated with T1D. Specifically, we searched for differences in the T-cell compartment between the healthy children and children with T1D, and for changes occurring during the first year after the diagnosis of T1D. We based our analysis on the single-cell transcriptomics by RNA sequencing (scRNAseq) including T-cell receptor sequencing (VDJ profiling) and flow cytometry analysis of T cells from the peripheral blood of our Czech cohort, which was complemented by the analysis of openly accessible data from other cohorts.

Blood of 30 donors with T1D was taken 4-9 (median 9) days after their diagnosis (time T0) and again at 359-423 (median 377) days after diagnosis (time T1). For comparison, we collected blood of 13 healthy donors (Fig. 1A). We characterized the donors by their age, sex, partial clinical remission status at T1, T1D-associated autoantibody levels, HLA haplotypes (Fig. 1B) as well as blood concentrations of glycated hemoglobin, random and fasting C-peptide, and leukocyte levels (Fig. S1A-C). The T1D patients showed higher overall blood lymphocytes and lower neutrophils at T0 in comparison to T1 and healthy donors (Fig. S1C).

**Figure 1.**
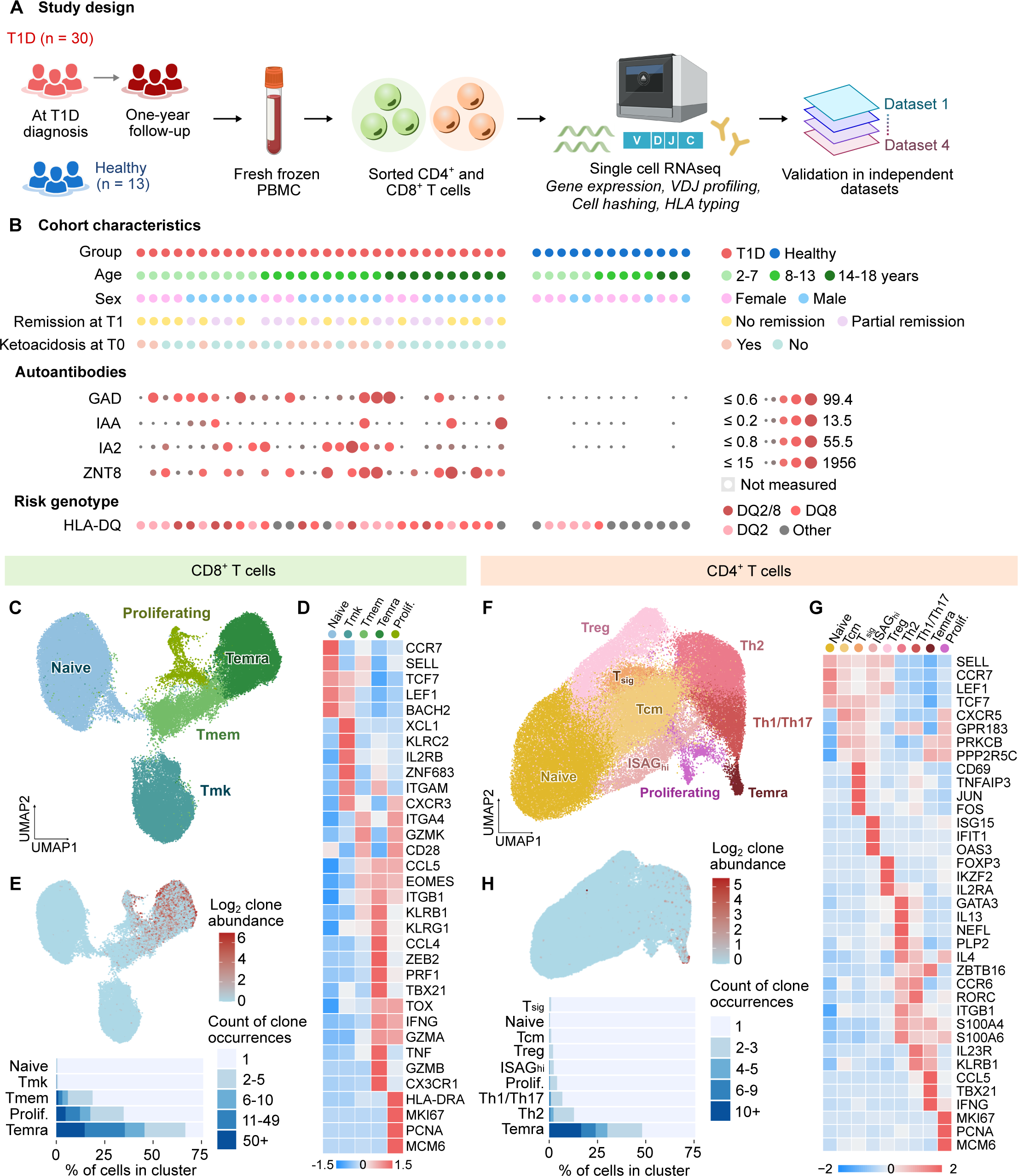
Blood T-cell atlas of children with T1D and healthy donors. A) Schematic representation of the experimental design. PBMCs from donors with T1D at diagnosis (n = 30) or at one-year follow-up (n = 29) and age- and gender-matched healthy donors (n = 13) were collected and fresh frozen. Sorted CD4^+^ and CD8^+^ lymphocytes were processed with scRNAseq (gene expression profiling and TCR repertoire profiling). Results were validated in independent cohorts from published data. B) Characteristics of the cohort. Each dot represents one participant of the study. In the upper part, colors represent the group, age, sex, remission state one year after diagnosis (T1) (remission is defined as insulin dose adjusted HbA1c <9) and diabetic ketoacidosis at diagnosis (T0) (ketoacidosis is defined as pH below 7.3 and/or bicarbonate <15 mEq/L). In the middle part, the levels of autoantibodies of the participants is indicated. In the bottom part, the presence of T1D risk HLA alleles inferred from the scRNAseq data is indicated. (C-E) CD8^+^ T cells were FACS-sorted from fresh-frozen PBMCs of children with T1D or healthy donors and processed with scRNAseq. Low quality cells, contaminating cells, unconventional CD8^+^ T cells and CD8^low^ NK cells were removed prior to this analysis. n = 63068 cells from 43 donors. C) UMAP projection of conventional CD8^+^ T-cell populations sorted from PBMCs of T1D patients and healthy donors and processed with scRNAseq. Louvain clusters were merged based on functional relevance and manually annotated based on marker genes. D) The heatmap of selected marker genes that characterize clusters presented in (C). Colors represent row-scaled z-score of average expression of a gene in a cluster. E) Analysis of TCR repertoires of CD8^+^ T-cell populations. In the upper part, the same UMAP projection as in (C) is shown, where cells are colored by their clonal expansion status. Expansion is calculated as the log2 counts of occurrence of the same TCR CDR3α and CDR3β sequences in the dataset. In the bottom part, the clonal expansion is quantified for clusters presented in (C). (F-H) CD4^+^ T cells were FACS-sorted from fresh-frozen PBMCs of T1D patients or healthy donors and processed with scRNAseq. Low quality cells, contaminating cells, and unconventional CD4^+^ T cells were removed prior to this analysis. n = 79,876 cells from 43 donors. F) UMAP projection of conventional CD4^+^ T-cell populations sorted from PBMCs of T1D patients and healthy donors and processed with scRNAseq. Louvain clusters were merged based on functional relevance and manually annotated based on marker genes. G) The heatmap of selected marker genes that characterize clusters presented in (F). Colors represent row-scaled z-score of average expression of a gene in a cluster. H) Analysis of TCR repertoires of CD4^+^ T-cell populations. In the upper part, the same UMAP projection as in (F) in which cells are colored by their clonal expansion status. Expansion is calculated as the log2 counts of occurrence of the same TCR CDR3α and CDR3β sequences in the dataset. In the bottom part, the clonal expansion is quantified for clusters presented in (F). T1D – Type 1 Diabetes Mellitus, T0 – timepoint at diagnosis, T1 – timepoint one year after diagnosis, Tmk – unconventional KLRC2^+^ memory T cells, Tmem – memory T cells, Tcm – central memory T cells, Treg – regulatory T cells, Tsig – T cells with signaling transduction signature, ISAGhi – T cells with interferon signaling signature, GAD – Glutamic Acid Decarboxylase, IAA – Insulin Autoantibodies, IA2 – Tyrosine Phosphatase-like Protein IA-2, ZNT8 – Zinc Transporter 8.

First, we performed an initial scRNAseq analysis of CD4^+^ and CD8^+^ lymphocytes from 16 samples, which revealed that the majority of T cells in our child donors had a naïve CD45RA^+^ CD45RO^-^ phenotype (Fig. S2A-D), determined by CD45 splicing-sensitive analysis tool IDEIS ^28^. To enrich the antigen-experienced T cells for the final scRNAseq analysis, we sorted the cells at a 1:5 fixed ratio of naïve vs. antigen-experienced cells for CD4^+^ and CD8α^+^ lymphocytes (Fig. S2E-H). We sequenced the cDNA libraries to an average depth of 2736 non-redundant transcripts per cell and processed the data using our standard pipeline including the removal of dead cells and doublets (Fig. S2I). Because the library preparation was performed in four batches, we removed the batch effects by data integration (Fig. S2J) ^29, 30^.

In the next step, we generated CD8^+^ and CD4^+^ lymphocyte atlases using the processed scRNAseq data. The CD8α^+^ lymphocyte compartment consisted of three clusters of CD8^low^ lymphocytes: NK T cells, semi-invariant MAIT cells and γδT cells, as well as conventional CD8^+^ αβT cells (Fig. S3A-D). Based on the typical markers, we identified that conventional CD8^+^ T cells contain naïve, memory (Tmem), unconventional KLRC2^+^ memory also previously described as KILR-like cells ^31, 32^, which we abbreviate as Tmk, terminally differentiated CD45RA^+^ (Temra), and proliferating subsets (Fig. 1C-D, Fig. S3E). Temra and to a lesser extent, proliferating and Tmem cells, but not Tmk cells, were clonally expanded (Fig. 1E). These subsets could be further subdivided into 41 smaller subsets (Fig. S3F).

The CD4^+^ lymphocyte compartment consisted of a subset of unconventional T cells including iNKT cells expressing their signature TCRα (*TRAV10*, *TRAJ18*; CDR3: CVVSDRGSTLGRLYF) and conventional T cells (Fig. S4A-B), which could be further divided to naïve, central memory (Tcm), Treg, Th1/Th17, Th2, Temra, ISAGhi ^33^, proliferating cells, and CD4^+^ T cells enriched for genes in signal transduction pathways (Tsig) (Fig. 1F-G, Fig. S4C-F). Temra cells were the only CD4^+^ T-cell subset showing a substantial level of clonal expansion (Fig. 1H). These subsets could be further split into 38 smaller subsets (Fig. S4G).

We compared the proportions of particular subsets using both frequentists (Fig. S5A-D) and Bayesian (Fig. S5E-F) statistics. These analyses did not reveal substantial differences between T1D children and healthy donors, with the exception of CD8^+^ Temra cells, which were enriched in the healthy donors in comparison to the T1D children (Fig. S5A, S5E). The Bayesian statistics revealed apparent differences in ISAGhi and Tsig subsets (Fig. S5F), which however, originated from only a limited number of outlier patients (Fig. S5C). Within the T1D group, the frequency of MAIT cells among CD8^+^ T cells and the frequency of Th2 cells among CD4^+^ T cells correlated with fasting C-peptide level, which we used as a proxy for residual β-cell activity (Fig. S5G-H). However, because of the prior enrichment for antigen-experienced T cells, the relative abundance of the particular subsets in the scRNAseq data does not correspond to the real frequency in the original sample.

### The onset of diabetes is associated with low effector T-cell signatures

Next, we compared gene expression between healthy donors and diabetic children at T0 and T1, separately in CD4^+^ and CD8^+^ T cells (Fig. S6A). Effector and cytotoxic genes (*GZMA, GZMB, GZMH, PRF1, GNLY, IFNG, TNF, CCL5, CCL4*) were downregulated, whereas genes associated with naïve, quiescent, and/or stem-like T cells (*LEF1, BACH2, IL7R, CXCR4, ZFP36L2*) were upregulated in the T1D children in comparison to the healthy controls, particularly at T0 and to lesser extent and somewhat variably at T1 (Fig. 2A-B). This bias towards naïve T-cell gene expression signature was more pronounced at T0 than at T1 (Fig. 2B). T cells from T1D T0 samples also showed higher expression of several regulators of intracellular activation signaling pathways (Fig. 2B), which might be associated with the quiescent state. We quantified the expression of genes previously described as upregulated or downregulated in naïve vs effector CD4^+^ or CD8^+^ T cells ^34, 35^ using Gene Set Enrichment Analysis (GSEA), which confirmed the enrichment of the preferential naïve-like signature in T cells from the T1D children and the enrichment of effector or memory specific genes in the healthy donors (Fig. 2C). In addition to the whole CD4^+^ and CD8^+^ T-cell compartments, a bias towards the effector gene expression signature was observed across multiple T-cell subsets (Fig. S6B).

**Figure 2.**
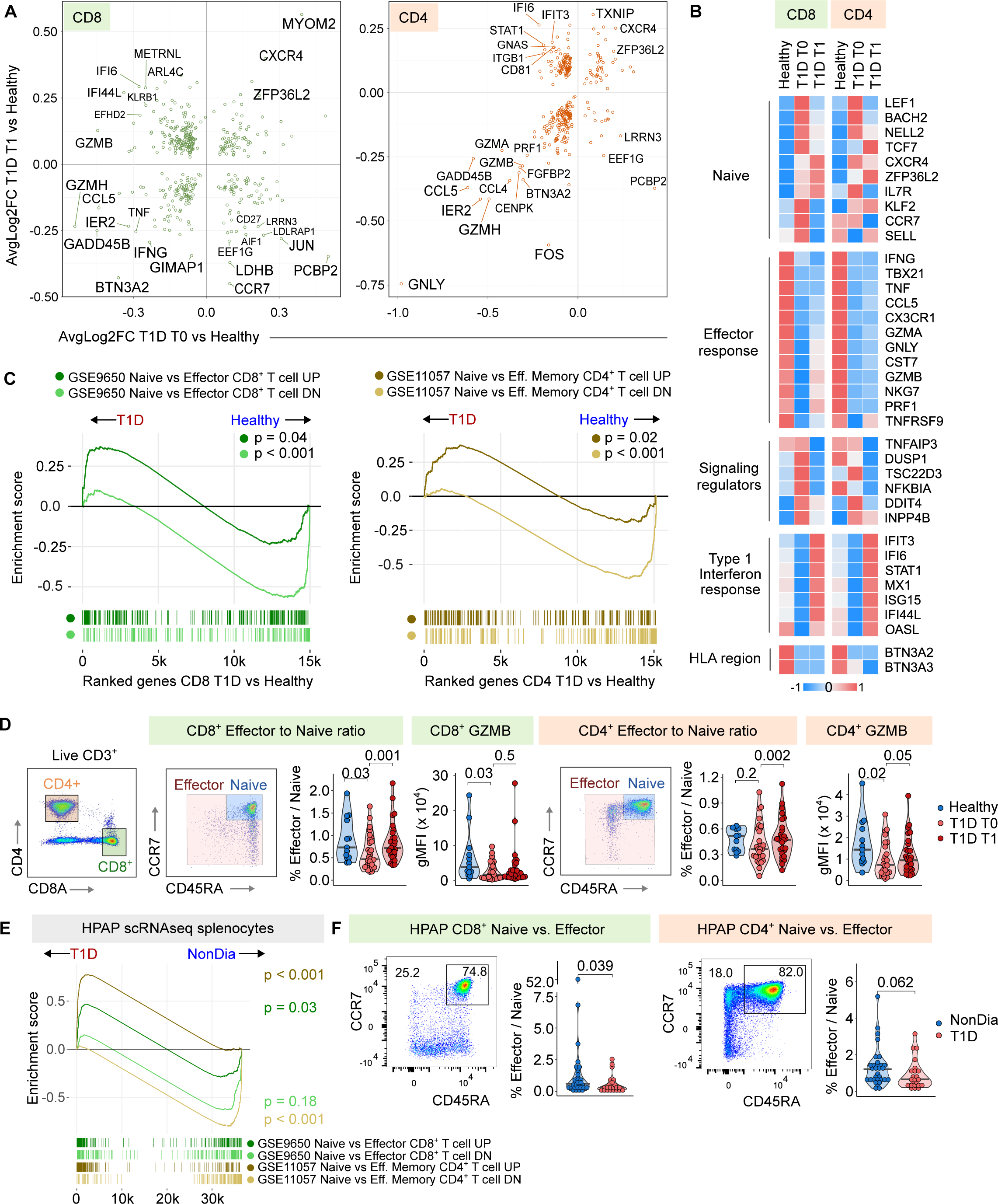
Differential gene expression in T-cells in children with T1D. A) Differentially expressed genes between T1D children and healthy donors at diagnosis (T0) and one year after diagnosis (T1) in CD8^+^ T cells (left) and CD4^+^ T cells (right) are visualized. The average log2 fold changes were calculated using the FindMarkers function from the Seurat package (Wilcoxon test). Genes with adjusted p-value < 0.05 (Bonferroni correction) were plotted and selected genes were manually labelled. B) The heatmap shows the relative expression of selected genes in healthy and T1D children at T0 and T1. Average expression of each gene per each patient was calculated using the function AverageExpression from the Seurat package. Plotted is z-score normalized average of donors in the healthy, T1D T0 and T1D T1 groups. C) Gene set enrichment analysis (GSEA) showing enrichment of genes identified as differentially expressed in previously published studies (GSE9650 ^35^, GSE11057 ^34^) in the CD8^+^ (left) or CD4^+^ T cells (right) from T1D children compared to healthy donors. D) Validation of scRNAseq results by flow cytometry. Effector and naïve T cells were gated from Live CD3^+^ CD8^+^ or CD4^+^ T cells. For CD8^+^ T cells (left) and CD4^+^ T cells (right), the following plots are shown (from left to right): Representative gating of naïve (CCR7^+^ CD45RA^+^) vs. effector populations, the effector to naïve ratio, and the geometric mean of the intensity of the GZMB signal in CD8^+^ or CD4^+^ T cells. Full gating is shown in Fig S2. P-value was calculated using two-tailed Mann-Whitney test between healthy donors and T1D T0, or two-tailed paired Mann-Whitney test between T1D donors at T0 and T1. Bar at median. n = 13 healthy donors, n = 30 T1D T0 donors, n = 29 T1D T1 donors. E) GSEA showing enrichment of genes identified as differentially expressed in previously published studies (GSE9650 ^35^, GSE11057 ^34^) in the splenocytes of T1D and non-diabetic donors from the HPAP dataset. F) Flow cytometry analysis of CD8^+^ and CD4^+^ T cells from the PBMCs of T1D and non-diabetic (NonDia) donors from the HPAP dataset. For CD8^+^ T cells (left) and CD4^+^ T cells (right), the following plots are shown (from left to right): Representative gating of naïve vs. effector populations and the effector to naïve ratio. Full gating is shown in Fig S7. P-value was calculated using two-tailed Mann-Whitney test. Bar at median. n = 32 non-dia donors, n = 23 T1D donors. T1D – Type 1 Diabetes Mellitus, T0 – timepoint at diagnosis, T1 – timepoint one year after diagnosis, GSEA – Gene set enrichment analysis

To check whether the downregulation of effector genes and upregulation of naïve genes in T1D children is associated with the disease severity, we carried out the comparison of differential expression between T1D children at T0 who suffered from ketoacidosis and those who did not, and between T1D who fulfilled or not the criterion of partial clinical remission (insulin dose adjusted HbA1c <9) (Fig. S6C-E). The naïve signature did not seem to be more pronounced in either of these subgroups.

For the interpretation of the previous results, it needs to be taken into account that the ratio of naïve and antigen-experienced T cells has been normalized for the scRNAseq. For this reason, we analyzed the pre-enriched samples by flow cytometry to reveal that both antigen-experienced CD8^+^ and CD4^+^ T cells were less abundant in patients than in healthy donors at T0, but not at T1 (Fig. 2D). Accordingly, the expression of a key cytotoxic protein Granzyme B (GZMB) is lower in the T1D patients in CD8^+^ and CD4^+^ T cells than in the healthy controls at both time points (Fig. 2D).

Overall, the data suggested that early T1D is accompanied with a more naïve state of the T-cell compartment than in the healthy controls. This naïve state is partially normalized within one year after the diagnosis and the associated introduction of the insulin therapy.

To extend our findings to unrelated T1D patient cohorts, we addressed the question of the naïve vs. effector signature in T1D patients in previously published data. Using the HPAP database ^27^, we re-analyzed scRNAseq data obtained from T cells from spleens of children and adult diabetic patients and non-diabetic donors. In this cohort, the effector signature genes were enriched in the non-diabetic controls and the naïve signature genes were enriched in the diabetic patients (Fig. 2E). In addition, we re-analyzed flow cytometry data from the HPAP and observed a reduction of effector CD8^+^ and CD4^+^ T cells in the blood of T1D patients (Fig 2F, Fig. S7A), consistent with the findings in our cohort.

Next, we collected transcriptomic data from previously published studies ^20, 23, 25, 36, 37^ (Supplemental Table 1) and re-analyzed it using the same pipeline. Among the genes that were identified across the datasets as the most upregulated in T1D children, we found genes connected to the naïve, stem-like T cells, and/or quiescent T-cell states such as *LEF1* ^38^*, FOXP1* ^39^, *BACH2* ^40^, *NELL2* ^41^, *IKZF1* ^42^ and a negative regulator PI3 kinase signaling *INPP4B* ^43^ (Fig. S7B). In contrast, genes associated with cytotoxicity and effector functions, such as *NKG7*, *GNLY*, *CCL4*, *CCL5*, *CX3CR1, TNFRSF4* (OX40)*, TNF,* and *TNFRSF9* (4-1BB) were upregulated in healthy donors (Fig. S7B). Altogether, our results showing a T-cell bias towards the naïve, stem-like, and quiescent phenotype in T1D patients are supported by the majority of the published datasets.

### BTN3A2 expression depends on HLA haplotypes

It was shown that *BTN3A2* and to a lower extent *BTN3A3,* encoding butyrophilin family members, are upregulated in T cells of T1D children in comparison to matched controls ^20^. However, we observed the opposite in our dataset, i.e., the higher expression of *BTN3A2* and *BTN3A3* in healthy donors than in T1D children (Fig. 2A-B). Since *BTN3A2* is a part of the extended MHC region ^44^, we hypothesized that its expression might depend on the particular HLA haplotype. First, we observed that our healthy donors generally followed the common distribution of HLA alleles in the Czech population ^45^ with the exception of overrepresented HLA-A*02:01, whereas most T1D children carried the respective susceptible alleles ^1^ (Fig. S8A). We used our data and publicly available data to estimate the role of the T1D status, particular haplotype, and study on *BTN3A2* expression. We observed that the study and HLA haplotypes, but not the T1D status, are the factors significantly influencing *BTN3A2* expression (Fig. S8B-C). This indicates that the selection of healthy controls (e.g., unmatched as in our study or partially matched as in ^20^) might have a great impact on the results concerning *BTN3A2* expression.

### Naïve-like regulatory T cells in T1D children

It has been proposed that the dysfunction of Treg cells might contribute to the loss of self-tolerance and the development of autoimmune diabetes in humans and animal models ^46, 47^. We identified four Treg subclusters labeled Treg1 to Treg4 (Fig. 3A). The gene expression analysis revealed that Treg1 represents Tregs with the most naïve phenotype, with high expression of naïve T-cell markers such as *CCR7, TCF7, SELL* and relatively low expression of Treg signature genes such as *FOXP3, CTLA4*, and *IL2RA* (Fig. 3B-D). However, the expression of these signature molecules in Treg1 cells was higher than in naïve T cells suggesting that they are *bona fide* Tregs (Fig. 3D). In contrast, the Treg4 subset represents Tregs with the most activated phenotype (Fig. 3B-D), which predispose them for potent regulatory functions ^48, 49^.

**Figure 3.**
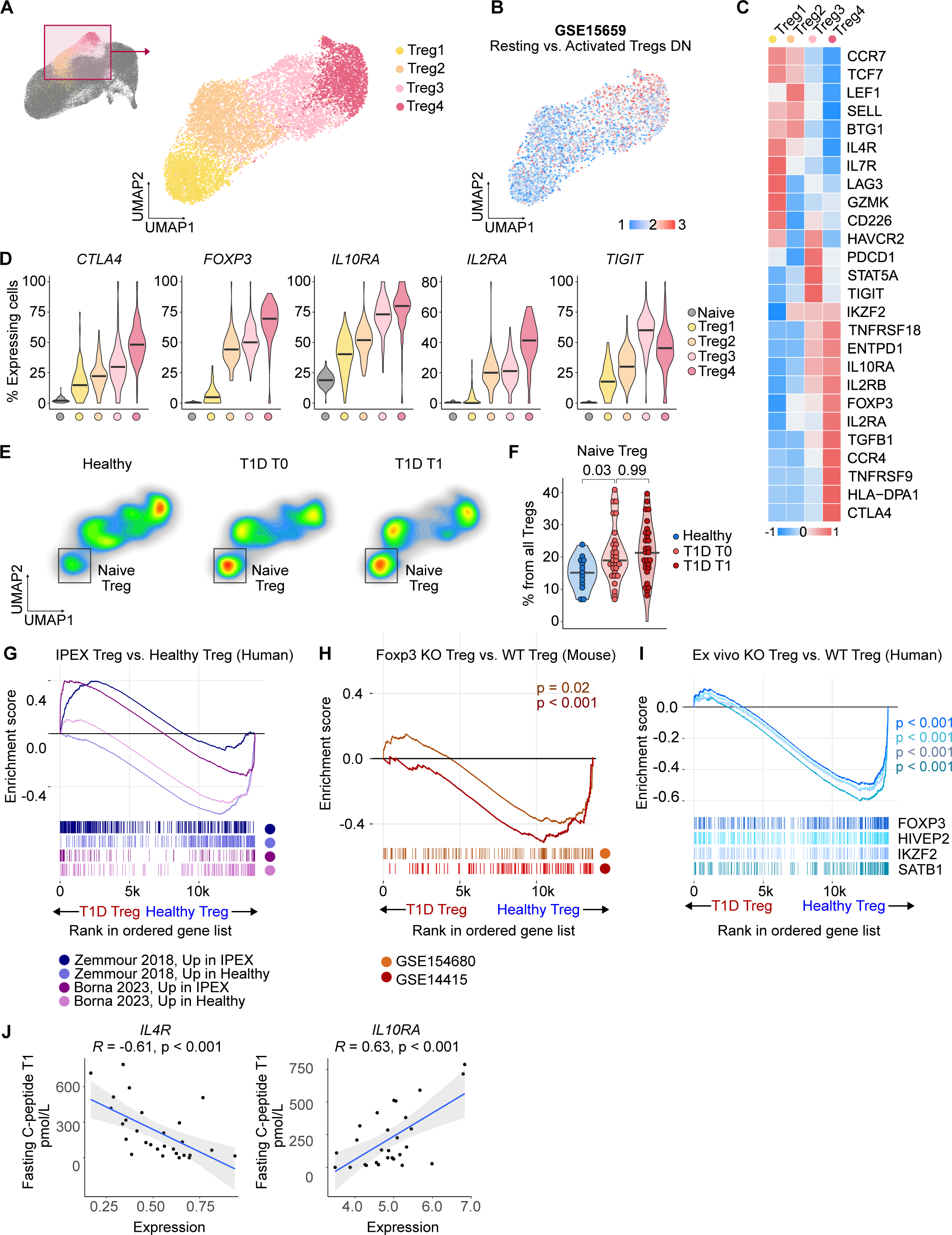
Regulatory T cells in children with T1D. A) Reclustering of Treg clusters from the UMAP projection shown in Fig. 1F. Selected clusters were extracted from the dataset and subjected to new normalization, scaling, integration, and dimensional reduction. On the right, the resulting UMAP projection is shown. n = 9,890 cells from 43 donors. Cells are colored by clusters. B) The same UMAP projection as in (A). The expression of the module of genes up-regulated in activated Tregs compared to resting Tregs from GSE15659 ^48^ is indicated. C) The heatmap of selected marker genes that characterize clusters presented in (A). Color represents row-scaled z-score of average expression of a gene in a cluster. D) Violin plots showing the percentage of cells from the clusters indicated in (A) that have non-zero expression of the selected gene within the specified cluster. The data are based on expression profiles from 87 samples. Bar at median. E) A density plot showing the difference between the abundance of specific Treg subtypes in samples coming from healthy donors or T1D patients at T0 and T1. F) A quantification of the percentage of cells in each sample representing the immature Tregs from total Tregs. Naïve Tregs were gated as shown on the UMAP projection in (E). P-value was calculated using two-tailed Mann-Whitney test between healthy donors and T1D T0, or two-tailed paired Mann-Whitney test between T1D children at T0 and T1. Bar at median. n = 13 healthy donors, n = 30 T1D T0 donors, n = 29 T1D T1 donors. G) Gene set enrichment analysis (GSEA) showing the enrichment of genes identified as upregulated or downregulated in patients with IPEX versus healthy donors ^54, 55^ in patients with T1D vs. healthy donors from our study. The ranked gene list represents the contrast between Tregs of children with T1D vs. healthy donors. H) GSEA showing the enrichment of genes identified as downregulated in *Foxp3* KO mice vs. control wild-type mice ^56^ ^57^ in children with T1D vs. healthy donors from our study. The ranked gene list represents the contrast between Tregs of patients with T1D vs. healthy donors. I) GSEA showing the enrichment of genes identified as downregulated in human Treg cells with CRISPR-mediated KO of FOXP3, HIVEP2, IKZF2, and SATB1 vs. normal human Treg cells ^58^ in children with T1D vs. healthy donors from our study. The ranked gene list represents the contrast between Tregs of children with T1D vs. healthy donors. J) Correlation of the levels of fasting C-peptide measured at T1 and the expression of *IL4R* (left) or *IL10RA* (right) within the whole Treg cluster (A) at T0. P-value was calculated using the Pearson’s correlation test. n = 28 T1D donors. T1D – Type 1 Diabetes Mellitus, T0 – timepoint at diagnosis, T1 – timepoint one year after diagnosis, GSEA – Gene set enrichment analysis, IPEX - Immunodysregulation polyendocrinopathy enteropathy X-linked syndrome

The naïve-like Treg subset was more abundant among Tregs in the diabetic children than in healthy donors (Fig. 3E-F, Fig. S9A) at T0 and T1. Moreover, the frequency of Treg1 cells among Tregs inversely correlated with residual insulin production measured as fasting C-peptide levels, whereas the frequency of Treg4 among Tregs correlated positively with fasting C-peptide levels (Fig. S9B), suggesting that the Treg maturation status might be involved in the disease severity. Treg1 cells expressed high levels of *CD226* and low levels of *TIGIT* (Fig. 3C), which both bind to common receptors CD112 and CD155, but have the opposite functions. CD226 is associated with Treg dysfunction, whereas TIGIT in Tregs is linked to the self-tolerance in the context of T1D ^50, 51, 52, 53^.

In the next step, we addressed whether Tregs from T1D ressemble functionally impaired Tregs from conditions in which their dysfunction is well established and characterized. First, we compared T1D Tregs from our cohort to Tregs from patients with a disease called *Immune dysregulation, polyendocrinopathy, enteropathy X-linked syndrome* (IPEX), which is caused by loss-of-function mutations in *FOXP3,* using a GSEA. Tregs from T1D patients upregulated genes specific for defective Treg-like cells observed in IPEX patients ^54, 55^ (Fig. 3G). We obtained the same results when we compared Tregs from the T1D patients to Treg-like cells in *Foxp3*^-/-^ mice ^56, 57^ (Fig. 3H) and *bona fide* human Tregs after their key transcriptional regulators *FOXP3*, *HIVEP2*, *IKZF2* (Helios), or *SATB1* were knocked-out ^58^(Fig. 3I). Moreover, we observed an inverse correlation between *IL4R* expression in Tregs and the fasting C-peptide levels (Fig. 3J), which points to a previously proposed inhibitory role of IL-4 signaling in Treg differentiation ^59, 60^. In contrast, the expression of *IL10RA* in Tregs correlated with fasting C-peptide levels, which highlights a possible role of IL-10 signaling for Treg-mediated tolerance in T1D (Fig. 3J). Overall, this evidence suggests that Tregs in T1D patients might be less functional than their counterparts in healthy controls.

To address our findings in independent cohorts, we extracted Treg clusters from three publicly available scRNAseq datasets containing T1D and healthy donors: a recent publication ^25^, HPAP dataset ^27^, and a reference dataset published by Parse Biosciences (henceforth ParseBio) (Fig. S9C-F). We reclustered these cells to obtain four clusters A-D, which roughly corresponded to our Treg1-4 clusters with A being the most naïve and D being the most mature (Fig. S9C-E). The cluster A, which showed high expression of naïve genes and *IL4R*, was overrepresented in Tregs from T1D donors in comparison to healthy controls, whereas the cluster D, which showed high expression of Treg effector genes, was overrepresented in healthy donors (Fig. S9F-G).

In the next step, we examined FOXP3^+^ Tregs using flow cytometry analysis of our cohort (Fig. S10A-B) and analogous data from the HPAP database (Fig. S10C-D). In both cases, there were slightly more FOXP3^+^ cells among CD4^+^ T cells in T1D children than in healthy donors. However, we did not see consistent differences in the expression of the naïve or effector signature genes between T1D children and healthy donors. A possible reason is the low expression of *FOXP3* in the naïve Treg subset (Fig. 3C-D, Fig. S9E), which makes it difficult to identify these cells by the conventional flow cytometry panels, which highly depend on FOXP3 detection.

### Low frequencies of effector-phenotype unconventional T cells in T1D patients

It has been proposed that a loss of unconventional CD3^+^ CD56^+^ T cells (named TR3-56) with regulatory properties might contribute to the development of T1D ^16, 61^. In our cohort, we observed that T1D children at T0, but not at T1, exhibited slightly lower expression of the TR3-56 signature than healthy donors, both in CD4^+^ and CD8^+^ T cells (Fig. 4A). We observed similar results in two unrelated cohorts with publicly available data (Fig. S11A).

**Figure 4.**
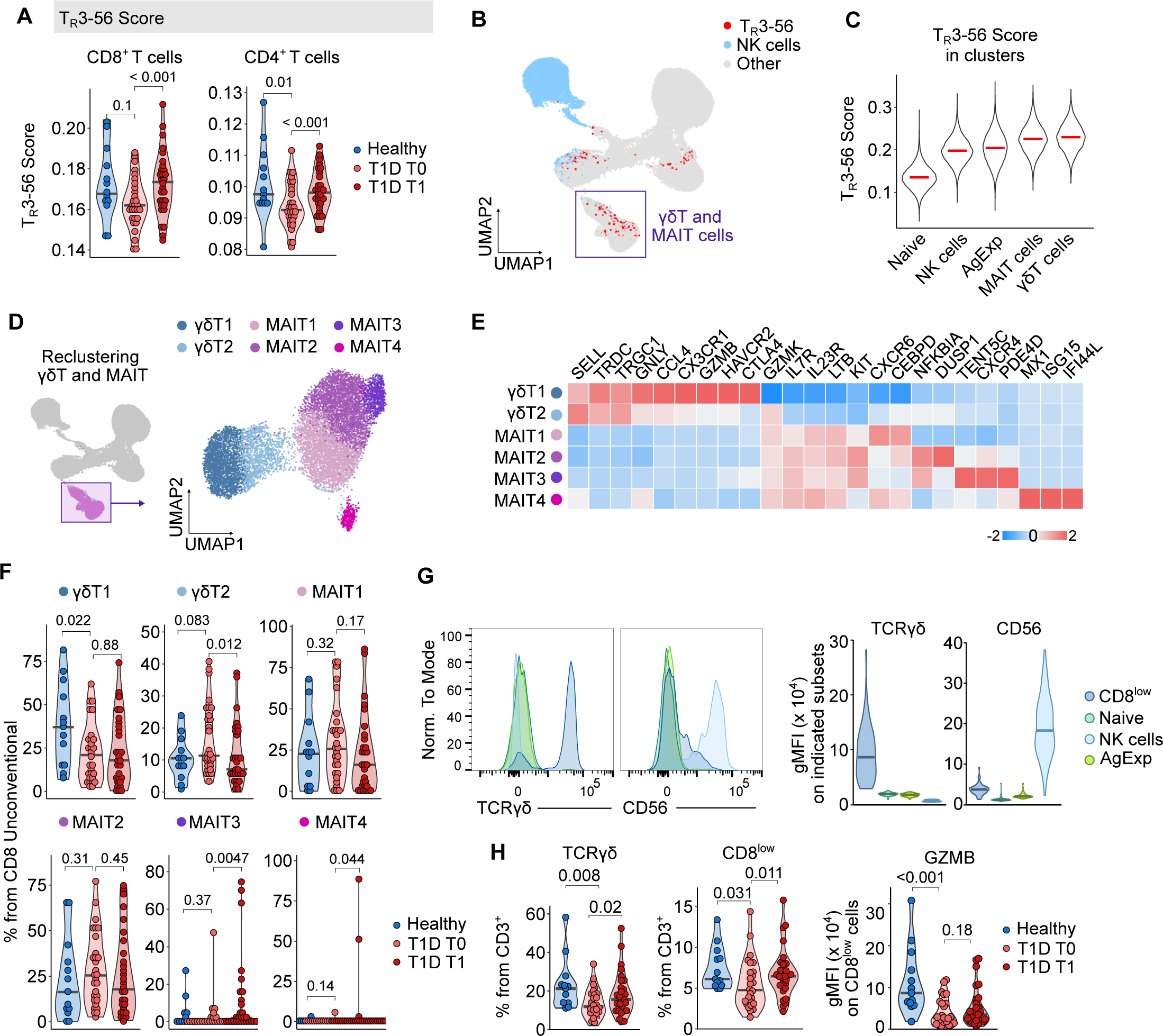
Unconventional CD8^+^ T cells in children with T1D. (A-C) Cells were annotated with a previously published RNA sequencing dataset of FACS-sorted TR3-56 (CD3^+^ CD56^-^) cells, NK cells, CD3^+^ CD56^-^, and CD8^+^ cells (GSE106082 ^16^ using the SingleR package. A) The quantification of the annotation scores of TR3-56 cells (i.e., similarity of the gene expression of a particular cell to that of previously published for TR3-56 cells) in CD8^+^ or CD4^+^ T cells in healthy donors and T1D children at T0 or T1. n = 13 healthy donors, n = 30 T1D T0 donors, n = 29 T1D T1 donors. B) The same UMAP projection as in Fig. S3A. Cells were annotated with a previously published dataset of FACS sorted TR3-56 (CD3^+^ CD56^-^) cells, NK cells, CD3^+^ CD56^-^, and CD8^+^ cells (GSE106082 ^16^) using the SingleR package. Cells annotated as NK and TR3-56 are indicated. C) Quantification of the annotation scores for different clusters. Violin plots show the annotation scores of TR3-56 cells. Medians are indicated. D) Reclustering of CD8^+^ γδT and MAIT cell clusters from the UMAP projection shown in Fig. S11B. Selected clusters (left) were extracted from the dataset and subjected to new normalization, scaling, integration, and dimensional reduction (right). n = 11,012 cells from 43 donors. Cells are color-coded by clusters. E) The heatmap of the relative expression of selected marker genes that characterize clusters presented in (D). Colors represent row-scaled z-score of average expression of a gene in a cluster. F) Percentage of cells in clusters shown in (D) from total Unconventional CD8^+^ T cells in healthy donors, and T1D children at T0 and T1. n = 13 healthy donors, n = 30 T1D T0 donors, n = 29 T1D T1 donors. (G-H) Flow cytometry analysis of CD8^+^ TR3-56 cells. Gating is shown in Fig. S11. G) The intensity of TCR γδ and CD56 was compared between naïve CD8^+^ T cells, non-naïve CD8^+^ T cells, NK cells, and CD8^low^ T cells. A representative histogram (left) and the quantification (right) are shown. Violin plots are based on intensity in 68 samples (n = 13 healthy donors, n = 26 T1D T0 donors, n = 29 T1D T1 donors). H) Percentage of γδT cells (left), or CD8^low^ T cells (middle) from CD3^+^ cells in healthy donors and T1D children at T0 and T1. The geometric mean of intensity of the GZMB in gated CD8^low^ cells (right). n = 13 healthy donors, n = 26 T1D T0 donors, n = 29 T1D T1 donors A, F, H - P-value was calculated using two-tailed Mann-Whitney test between healthy and T1D T0 donors, or two-tailed paired Mann-Whitney test between T1D donors at T0 and T1. Bar at median.

In the next step, we aimed to identify TR3-56 cells in our scRNAseq data, starting with the CD8^+^ T cells atlas, consisting of naïve and non-naïve conventional T, NK, γδT cells, and MAIT cells (Fig. 4B-C, Fig. S3A, Fig. S11B). From these cell types, it was mostly MAIT and γδT cells that coexpressed CD3 and CD56 (Fig. 4B-C, Fig. S11C). Indeed, the respective cell clusters contained most cells annotated as TR3-56 based on their gene expression signature ^16^(Fig. S11D). This suggests that TR3-56 cells at least partially overlap with MAIT and γδT cells.

Re-clustering of the unconventional T cells identified two clusters of γδT cells and four clusters of MAIT cells (Fig. 4D-E). We observed that the γδT1 cluster was enriched in healthy donors in comparison to T1D patients at T0 and T1 (Fig. 4F). This cluster corresponded to effector-phenotype γδT cells (Fig. 4E), suggesting that the state of γδT cells in T1D essentially phenocopies conventional T cells (Fig. 2A-D). In contrast, γδT2 cells, enriched in some T1D patients (Fig. 4F), expressed lower levels of effector genes (Fig. 4E). Accordingly, the total unconventional CD8^+^ T cells from T1D patients express lower levels of cytotoxic effector genes such as *GZMB*, *GZMH*, *GNLY, CCL5* (Fig. S11E).

Using flow cytometry, we observed that CD8^low^ T cells are enriched for γδTCR^+^ and CD56^+^ cells (Fig. 4G, Fig. S11E). Thus, gating for CD8^low^ cells can be used as a proxy for CD8^+^ γδT cells or CD8^+^ CD3^+^ CD56^+^ T cells in flow cytometry panels which lack these markers. In our flow cytometry analysis, we found that percentages of γδT cells and CD8^low^ T cells among CD3^+^ cells were low in T1D patients at T0, which was partially reverted at T1 (Fig. 4H, S11F). Moreover, a higher percentage of the CD8^low^ T cells expressed *GZMB* in healthy donors than in T1D patients, essentially confirming the scRNAseq data (Fig. 4H). A higher percentage of CD8^low^ T cells among CD3^+^ T cells from healthy donors compared to T1D donors was observed also in the HPAP flow cytometry data (Fig. S11G-H).

In the CD4^+^ T-cell compartment, the Temra cluster contained most TR3-56 phenotype cells (Fig. 5A-C) and had the gene expression profile most similar to TR3-56 cells (Fig. 5D-E). Re-clustering of the Temra cluster revealed four Temra subclusters and a contaminating NK cluster (Fig. 5F). The Temra2 cluster, corresponding to the cells with the strongest cytotoxic phenotype and the highest expression of *GZMB* (Fig. 5G), was more abundant in healthy donors than in T1D patients at T0 (Fig. 5H).

**Figure 5.**
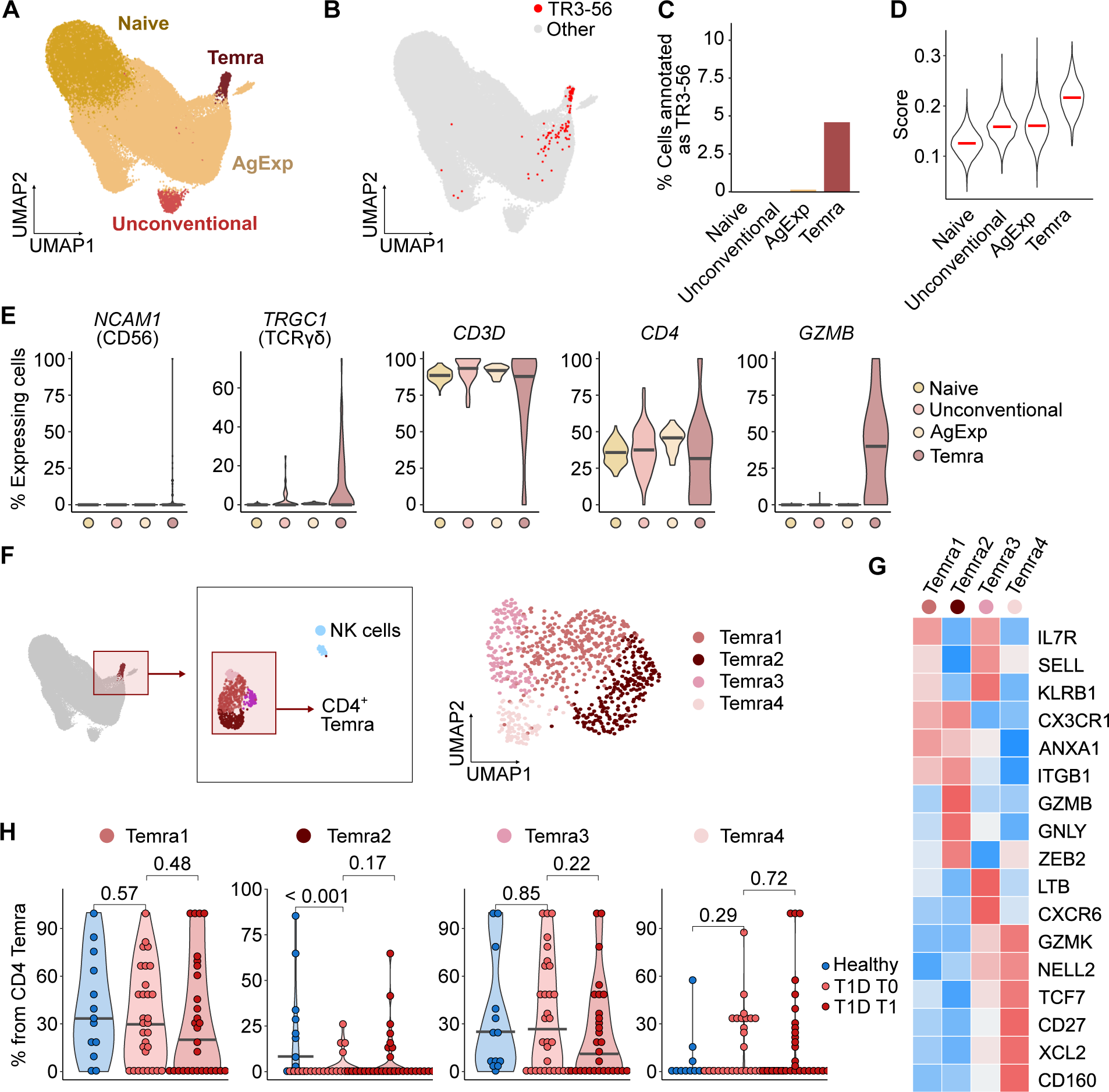
Unconventional CD4^+^ T cells in children with T1D. A) The UMAP projection of CD4^+^ T cells shows groups of cells used for the analysis of CD4^+^ subsets. Louvain clusters were merged based on functional relevance. n = 79,876 cells from 43 donors. B) Cells were annotated with a previously published dataset of FACS sorted TR3-56 (CD3^+^ CD56^-^) cells, NK cells, CD3^+^ CD56^-^, and CD8^+^ cells (GSE106082 ^16^) using the SingleR package. Cells annotated as TR3-56 are indicated. C) Percentage of cells in each cluster annotated as TR3-56 cells. D) Violin plots show the annotation scores of TR3-56 cells (i.e., similarity of the gene expression of a particular cell to that of a TR3-56 cell) in the indicated clusters. Medians are indicated. E) Violin plots show the percentage of cells from the indicated clusters as in (A) that have non-zero expression of the indicated genes within the specified cluster. Data is based on expression profiles from 87 samples. Bar at median. F) Reclustering of the CD4^+^ Temra cluster from the UMAP projection shown in (A). CD4^+^ Temra cells were extracted from the dataset (left) and subjected to new normalization, scaling, integration, and dimensional reduction (right). Contaminating NK cells, which formed a separate cluster after re-clustering, were removed. n = 815 cells from 43 donors. G) The heatmap shows the relative expression of selected marker genes that characterize clusters presented in (F). Color represents row-scaled z-score of average expression of a gene in a cluster. H) Percentage of cells in subclusters shown in (F) from total CD4^+^ Temra cells in healthy and T1D donors at T0 and T1. P-value was calculated using two-tailed Mann-Whitney test between healthy and T1D T0 donors, or two-tailed paired Mann-Whitney test between T1D donors at T0 and T1. Bar at median.

### The analysis of T-cell receptor repertoire in T1D

Together with the single-cell transcriptomics, we analyzed T-cell receptor sequences in T cells from the T1D children and healthy donors. We observed only minimal clonal overlap between individual donors regardless of their disease status (Fig. 6A). However, there was a significant overlap of TCRα and TCRβ sequences in the same T1D children at T0 and T1 (Fig. 6A), indicating the persistence of these clones in the blood during the one year period. We did not identify any T-cell clones previously associated with T1D diabetes. The usage of variable, diversity, and joint segments did not significantly differ between the T1D patients and healthy donors (Fig. S12-13).

**Figure 6.**
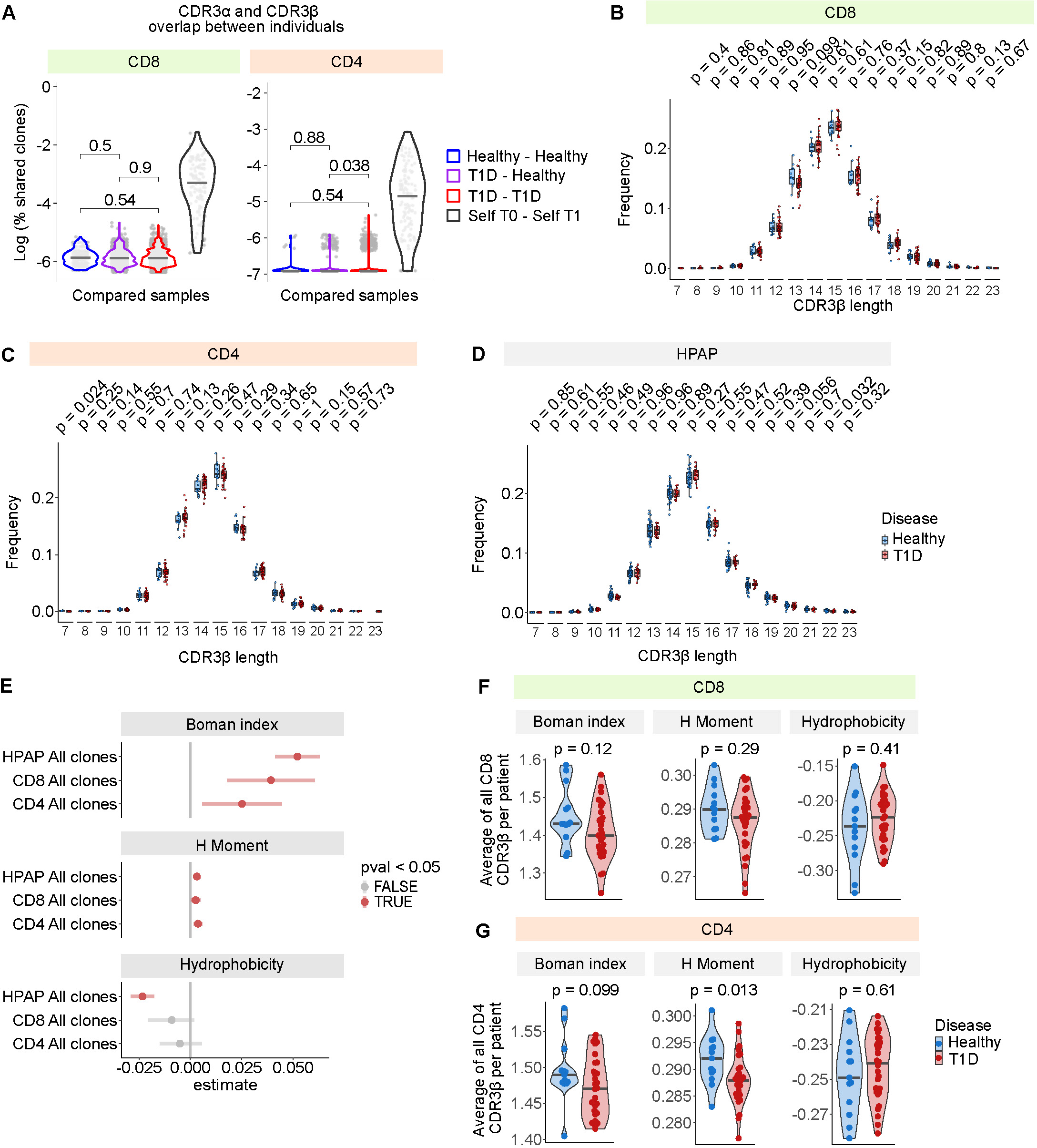
TCR repertoires in children with T1D. (A-C) TCR repertoires of the CD4^+^ and CD8^+^ T cells from patients with T1D at T0 (n = 30) or T1 (n = 29) and healthy donors (n = 13) were analyzed using 10x Immune profiling. (A) For each patient, unique amino-acid sequences of the CDR3 region of TCRα chain and TCRβ chain were extracted and used for the overlap analysis. For each participant, the percentage of overlapping sequences with all other participants was calculated and quantified. We quantified the overlaps between healthy donors with other healthy donors (blue color), healthy donor with a T1D donor (violet color) or two T1D donors (red color). The self T1 – self T0 comparison represents the overlap between clones in the same T1D patient at the two timepoints. The analysis was calculated for CDR3 sequences from CD8^+^ conventional cells after the exclusion of MAIT cells, NK cells, and γδT cells clusters in case of CD8^+^ T cells, or from conventional CD4^+^ T cells after the exclusion of iNKT cells in case of CD4^+^ T cells. P-value was calculated using two-tailed Mann-Whitney test. (B-D) For each patient, unique amino-acid sequences of the CDR3β regions were extracted and used for the length analysis. The lengths of the sequences were quantified as the frequency of each length in a particular donor and visualized in T1D patients and healthy donors. The analysis was calculated for CDR3 sequences from conventional CD8^+^ cells after the exclusion of MAIT cells, NK cells, and γδT cells clusters (B), from conventional CD4^+^ T cells after the exclusion of iNKT cells (C), or from mixed CD8^+^ and CD4^+^ T cells from the spleens of non-diabetic and T1D donors in the HPAP dataset (D). P-values were calculated using two-tailed Mann-Whitney test without a correction for multiple comparisons. (E-G) For each donor, unique amino-acid sequences of the CDR3β regions were extracted and used for the analysis of biochemical properties using the package Peptides ^119^. For each of the biochemical features (Boman index, H Moment, Hydrophobicity), the indicated indexes were calculated for all unique CDR3β regions from conventional cells CD8^+^ after the exclusion of MAIT cells, NK cells, and γδT cells clusters (n = 30 T1D patients, n=13 healthy donors), from conventional CD4^+^ T cells after the exclusion of iNKT cells (n = 30 T1D patients, n=13 healthy donors), and from mixed CD8^+^ and CD4^+^ T cells from the spleens in the HPAP dataset (n = 17 T1D, n = 39 non-diabetic). (E) All clonotypes were pooled within a group of donors and the contrast between T1D and non-diabetic donors is shown for the indicated groups (our CD4^+^ T cells, our CD8^+^ T cells, HPAP T cells). P-value was calculated using two-tailed Mann-Whitney test comparing T1D patients vs healthy donors. The estimate represents the difference of the location, i.e. the median of the difference between a CDR3 β sequence from T1D donor and a CDR3β sequence from a healthy donor. Lines represent the 95 % confidence interval of the difference of the location between T1D patients and healthy donors. (F-G) For each of the biochemical features, the values for all the unique CDR3β regions were averaged for each donor in our cohort. The differences in these average values between T1D and healthy donors is shown for CD8^+^ T cells (F) and CD4^+^ T cells (G). P-value was calculated using two-tailed Mann-Whitney test. Bar at median. T1D – Type 1 Diabetes Mellitus, HPAP – Human Pancreas Analysis Program

It has been reported previously, that the CDR3 segments of TCR β-chains in T1D patients are generally shorter than in healthy individuals ^62^. However, we did not observe any differences in the CDR3β length in CD4^+^ or CD8^+^ T cells in our cohort or in TCR repertoire profiling data from the HPAP database (Fig. 6B-D).

Finally, we analyzed the biochemical properties of the CDR3β TCR sequences in our cohort and in the HPAP cohort. We observed that the CDR3β sequences from T1D patients showed significantly higher Boman coefficient and H Moment and a slightly lower Hydrophobicity index (Fig. 6E). This effect was apparent also on the average values per patient (Fig. 6F-G), but with a relatively low significance. A large cohort is required for eventual confirmation of this observation.

## Discussion

T1D is an incurable autoimmune disease that is often diagnosed in childhood, unlike many other autoimmune diseases. Genetic factors, such as susceptible HLA haplotypes and polymorphisms in immune regulating genes, as well as environmental factors contribute to the onset of the disease. However, it is still unclear how T1D is triggered. Moreover, the pathological mechanisms by which dysregulated immune actions lead to the destruction of insulin-producing β-cells remain unclear. Evidence such as the involvement of HLA loci ^1^ and requirements of T cells for autoimmune diabetes in animal models ^63^ suggested that T cells play a key role in T1D. Accordingly, T-cell depletion by anti-CD3 monoclonal antibody Tepluzimab is approved by United States Food and Drug Administration for the application to children with pre-clinical T1D to delay the onset of the clinical stage ^64, 65^. Recently, beneficial effects of this treatment were observed even when administred to individuals with clinical T1D ^66^.

In this study, we analyzed peripheral blood T cells of newly diagnosed T1D children and healthy donors using scRNAseq and flow cytometry. We have chosen this early time point to capture the state of the immune system around the disease onset. To compare how the immune system changes in time, we analyzed the same children at one year after the diagnosis. One of our major findings was that the T cells from the T1D patients showed a reduced effector phenotype in comparison to controls. This was apparent in the conventional CD4^+^ and CD8^+^ T cells, but also in unconventional subsets such as γδT cells. The cytotoxic genes such as *GZMB* were downregulated in T cells from T1D patients across various subsets, which is counterintuitive given the fact that direct T-cell cytotoxicity against pancreatic β-cells has been proposed as a T1D-inducing mechanism ^67, 68, 69^. The down-regulation of the effector genes becomes less pronounced at one year after diagnosis, but still apparent in comparison to healthy donors. These results were supported by our analysis of the splenic scRNAseq and blood flow cytometry data available at the HPAP ^27^ and by results of our reanalyses of the majority previously published RNAseq datasets. Accordingly, previously published flow cytometry analysis observed higher frequencies of naïve T cells and lower frequencies of terminally differentiated Temra cells in adults at the onset of T1D in comparison to healthy controls ^70^. Experiments on non-obese diabetic (NOD) mice showed that lymph node resident stem-like self-reactive pancreatic-specific CD8^+^ T cells were more potent in the T1D triggering than their effector counterparts ^71^. The stem-like T cells migrated to pancreatic islets and generated the cytotoxic effectors *in situ*. Accordingly, it has been shown that pancreatic β cell-specific CD8^+^ T cells retain stemness-associated epigenetic signature in the blood of T1D patients ^72^. Overall, the reduced frequency of effector T cells and cytotoxic T-cell signatures in T1D patients does not contradict these recent advances in the field.

The cause of this naïve-phenotype bias of the T-cell compartment in the T1D patients is unclear. One possibility is that it is the consequence of the metabolic imbalance induced by the disease. However, we did not see particularly low expression of the effector genes in patients with ketoacidosis, documenting that it does not correlate with the severity of the metabolic disorder. Another possible explanation is that the insulin deficiency directly affects the state of the T cell compartment via hypostimulation of the insulin receptor signaling pathway, which was previously shown to promote T-cell effector function in mice in an intrinsic manner ^73^. However, we observed reduced expression of the cytotoxic genes in patients one year after their diagnosis, albeit less pronounced than upon diagnosis, and in adults with T1D from the HPAP database, which were already compensated by the exogenous insulin intake. Thus, we find it more probable that the phenomenon of low effector and cytotoxic profile in T cells precedes or coincides with the onset of clinical T1D. This scenario is in line with the hygiene hypothesis, which explains the current outbreak of autoimmune diseases in high-income countries by a reduced exposure of individuals to infectious agents leading to immaturity of the immune system, which results in the loss of self-tolerance ^10, 74, 75^. The evidence for the hygiene hypothesis in T1D comes from experiments on NOD mice, which show higher incidence of diabetes in germ-free conditions than in standard housing facilities ^76^, and from population studies ^77, 78^. An apparently contradictory hypothesis postulates that T1D is triggered by viruses, such as enteroviruses ^7, 8, 79^. However, these two phenomena can actually be coupled as it is possible that individuals with an immature immune system and/or those being infected at older age with the hypothetical triggering virus might be at a particularly high risk of by-stander T1D development. This scenario goes along with the hypothesis that the immune system is shaped by the environment in a relatively short window in the early life ^80^.

Tregs are crucial for maintaining peripheral tolerance. For that reason, the lack or dysfunction of Tregs has been studied as a potential cause in many autoimmune diseases ^81^. In the T1D research, controversial data have been published, showing reduced ^70, 82^, normal ^83, 84, 85, 86, 87, 88, 89^, or elevated ^90, 91, 92^ FOXP3^+^ Treg frequencies in the peripheral blood in individuals with T1D compared to healthy controls. Our and HPAP flow cytometry data showed slightly elevated Tregs among CD4^+^ T cells in T1D. These discrepancies might be at least partially caused by differences in the age or time after T1D onset in the particular cohorts. Accordingly, it has been shown that the frequencies of Tregs increase within the first year after T1D diagnosis ^93^.

Partial dysfunction of Tregs as well as their impaired maturation phenotype in individuals with T1D have also been proposed ^25, 84, 85, 90, 91, 94, 95, 96^. Despite the controversies and open questions surrounding the role of Tregs in T1D, Treg-targeted therapies have been used to treat T1D in preclinical models ^97^ and are being tested in clinical trials in T1D patients ^98, 99, 100^. In this study, we observed that a relatively high proportion of Tregs from T1D patients show low expression of key Treg effector genes such as *IL2RA*, *FOXP3*, *CTLA4*, *TGFB1* and high expression of *CD226, IL4R* and *GZMK*, which are connected with Treg dysfunction ^50, 51, 59, 60^ or T-cell senescence ^101^. Moreover, the whole Treg compartment was slightly altered in T1D patients, as it was enriched for the gene signature specific for dysfunctional *FOXP3*-deficient Treg-like cells such as those present in IPEX patients or *FOXP3* knock-out Tregs ^54, 55, 56, 57, 58^. T1D is one of the typical autoimmune symptoms in IPEX ^102^. In contrast to the pleiotropic IPEX syndrome, the patients in our cohort developed only T1D and were still self-tolerant to most tissues, indicating that their Treg cells were still largely functional. However, their Treg compartment might be prone to failure in a specific context such as suppressing an autoimmune response towards the pancreatic β-cells ^103^. Accordingly, the progressive loss of a specific subset of effector KLRG1^+^ ICOS^+^ Tregs is associated with the T1D development in NOD mice ^104^. We speculate that the bias of the Tregs in T1D patients towards the less effector phenotype and their similarities to genetically dysfunctional Tregs might contribute to T1D development.

A population of CD3^+^ and CD56^+^ coexpressing lymphocytes, called TR3-56, was proposed to be a regulatory subset preventing diabetes ^16^. This subset has never been identified by single cell transcriptomics. We did not see CD3 and CD56 co-expressing T cells as a standalone subset, but rather as a heterogeneous group of cells overlapping with unconventional subsets such as CD8^low^ γδT and MAIT cells, and CD4^+^ Temra cells. We observed generally reduced frequencies of these subsets in T1D patients in comparison to healthy controls at the time of their diagnosis. Low frequencies of γδT in T1D patients upon diagnosis and one year later has been reported previously ^105^. The same study showed a drop in CD56^+^ γδT cells at the time of T1D diagnosis, which was reverted after one year, which aligns with our data ^105^. Moreover, these unconventional cells showed generally more naïve and less effector phenotype in T1D patients at both time points in comparison to healthy controls in our cohort. It is unclear if these unconventional T cells are involved in the self-tolerance and T1D prevention or if their less cytotoxic phenotype only corresponds to the overall state of the T cells compartment in T1D children.

We used the data from our cohort to address previously proposed differences between T1D patients and healthy donors. Whereas a previous study observed elevated expression of butyrinophillin-encoding genes BTN3A2 and BTN3A3 in T1D patients ^20^, we observed the opposite. Our analysis of data generated by multiple studies showed that the expression of BTN3A2, residing in the extended MHC region, depends on particular HLA haplotypes. Whereas our healthy controls were not HLA-matched to the T1D patients, the previous study used partially matched controls ^20^, which might explain the different observations.

Our analysis of the TCR repertoires revealed clonal expansion in the antigen-experienced subsets and showed that same clones can be detected in the T1D patients one year after analysis. However, we could not detect any public clones present in different donors. Our cohort as well as data from the HPAP did not show changes in the CDR3 length between T1D patients and healthy donors, which were observed previously ^62^. We detected small differences in the average biophysical parameters of the TCRs between T1D patients and healthy controls, such as Boman index, H Moment, and hydrophobicity. However, additional studies are required to validate these findings.

Our analysis is the first comprehensive analysis of T cells from newly diagnosed T1D and the same patients at one year after the diagnosis using single cell transcriptomics on a medium-sized cohort. It indicated differences between the patients and healthy donors at the time of diagnosis, some of which persist to the one-year time point post diagnosis. Our findings indicate an overt naïve phenotype in the T-cell compartment, potential decreased functionality in the Treg compartment, and alterations in unconventional T-cell subsets. These findings were largely backed-up by our flow cytometry analysis and/or analyses of data published by others, when possible. However, our experimental approach still needs to be taken as an exploratory study on a size-limited cohort. Further studies addressing these conclusions are needed.

## Methods

### Ethical consent

The study was approved by institutional Ethics Committee (EK-819/20) of the Motol University Hospital. Written informed consent was obtained from all the participants and their legal guardians.

### Collection of clinical metadata

The samples were collected between February 2021 and February 2023 at a setting of a tertiary center for pediatric diabetes (Motol University Hospital, Prague, Czechia). After the consent of parents/caregivers and the study participants was granted, two study samples were obtained from each participant. The first sample was obtained at a median of 9 (4-24) days after the clinical diagnosis of T1D, the second sample at a median of 377 (359-423) days after diagnosis. At both time-points, fasting C-peptide, HbA1c, and the standard blood count with the differential count were performed. At T0, the measurements of the T1D specific autoantibodies (a-GAD, IAA, a-IA2, a-ZnT8) were performed. Data on diabetic ketoacidosis status at first contact (pH below 7.3 and/or bicarbonate <15) were obtained from individual medical records. To assess partial clinical remission at T1, we employed the insulin dose adjusted HbA1c (IDAA1c) as a marker. IDAA1c was calculated as HbA1c (%) plus [4 times insulin dose (units per kilogram per 24 h)]. IDAA1c <9 was considered as the presence of partial clinical remission ^106^.

The samples were obtained from 30 children with T1D at T0 and from 29 of these children at T1. One T1D participant withdrew from the study before the one-year follow-up visit. Their sample was included in all analysis except for paired statistical testing. A cohort of 13 healthy controls was also enrolled. The parents/caregivers and the participants signed a written consent.

### Processing of blood samples

Three to ten mL of peripheral blood were collected into EDTA-coated tubes, kept on ice, and transferred from Motol University Hospital to the Institute of Molecular Genetics within two hours. PBMCs were separated using Ficoll-Paque (GE Healthcare) and immediately frozen following a cryopreservation protocol (10x Genomics). Briefly, 4 ml of Ficoll-Paque was overlaid with blood and centrifuged at 400g for 30 minutes at room temperature (brake set to one). The mononuclear cell layer was washed in PBS and resuspended in RPMI medium containing 40% FBS. After adding an equal volume of freezing medium (RPMI, 40% FBS, and 30% DMSO), two to five aliquots of PBMCs were frozen and stored in liquid nitrogen. PBMCs were gently thawed by slow, sequential dilution in RPMI medium containing 10% FBS. To minimize cell stress, wide-bore tips were used during the thawing process.

### ScRNAseq experimental design

ScRNAseq was performed in two different experiments, hereafter referred to as the initial experiment and the final experiment.

### Initial scRNAseq experiment

The initial experiment was performed in two batches: batch 1 – CD8^+^ T cells from 6 patients and 2 controls in one well, batch 2 – CD4^+^ T cells from 6 patients and 3 controls, CD8^+^ T cells from 1 control in one well, and CD4^+^ and CD8^+^ cells from 6 patients and 3 controls in two wells. Cells were incubated on ice for 5 minutes with Human TrueStain FcX (BioLegend #422301) and for additional 30 minutes with anti-human CD4 and CD8 antibodies (CD8 APC, LT8, Exbio #1A-817-T100; CD4 AF700, MEM-241, Exbio # A7-539-T100) and one of the hashtag antibodies (TotalSeq™-C0251 anti-human Hashtag 1 Antibody, LNH-94 2M2, BioLegend #394661; TotalSeq™-C0252 anti-human Hashtag 2 Antibody, LNH-94 2M2, BioLegend #394663; Totalzeq™-C0253 anti-human Hashtag 3 Antibody, LNH-94 2M2, BioLegend #394665; TotalSeq™-C0254 anti-human Hashtag 4 Antibody, LNH-94 2M2, BioLegend #394667; TotalSeq™-C0255anti-human Hashtag 5 Antibody, LNH-94 2M2, BioLegend #394669; TotalSeq™-C0256 anti-human Hashtag 6 Antibody, LNH-94 2M2, BioLegend #394671; TotalSeq™-C0257 anti-human Hashtag 7 Antibody, LNH-94 2M2, BioLegend #394673; TotalSeq™-C0258 anti-human Hashtag 8 Antibody, LNH-94 2M2, BioLegend #394675; TotalSeq™-C0259 anti-human Hashtag 9 Antibody, LNH-94 2M2, BioLegend #394677; TotalSeq™-C0260 anti-human Hashtag 10 Antibody, LNH-94 2M2, BioLegend #394679) and with Hoechst 33258 for viability right before sorting. From each sample 10,000 CD4^+^ cells and 10,000 CD8^+^ cells were sorted. All samples were collected to the same collection tube, washed with PBS/0.05% BSA and counted using the TC20 Automated Cell Counter (#1450102, Bio-Rad). The viability of the cells before loading was higher than 85%.

### Final scRNAseq experiment

The final experiment was performed in four batches: batch 1 – CD4^+^ and CD8^+^ T cells from 8 patients T0, 8 patients T1, 4 controls in 4 wells, batch 2 - CD4^+^ and CD8^+^ T cells from 8 patients T0, 8 patients T1, 4 controls in 4 wells, batch 3 - 8 patients T0, 7 patients T1, 3 controls in 4 wells, batch 4 - 6 patients T0, 6 patients T1, 4 controls in 4 wells. Cells from the same patient from time 0 and time 1 were always processed in the same well to minimize batch effect. Cells were incubated on ice for 5 minutes with Human TrueStain FcX (BioLegend #422301) and for additional 30 minutes with anti-human CD4, CD8, CD45RA and CCR7 antibodies (CD4 APC, MEM-241, Exbio #1A-359-T100; CD8 PE, MEM-31, Exbio #1P-207-T025; CD45RA FITC, MEM-56, Exbio #1F-223-T100; CCR7 PeCy7, G043H7, BioLegend #353226) and one of the hashtag antibodies (the same as in the initial experiment) and with Hoechst 33258 for viability right before sorting. Non-naïve cells were enriched in the final sample as follows: from each sample 1 000 naïve (CCR7^+^ CD45RA^+^) and 5 000 antigen-experienced cells were sorted into two tubes, one for CD4^+^ T cells and one for CD8^+^ T cells. All samples were collected to the same two collection tubes, washed with PBS/0.05% BSA and counted using the TC20 Automated Cell Counter (#1450102, Bio-Rad). The viability of the cells before loading was higher than 85%. One control sample was removed from the analysis because of bad quality of data at sort.

Cells from both cohorts were loaded onto a 10x Chromium machine (10x Genomics) aiming for a the yield of 1500 cells per sample. cDNA libraries were prepared using the Feature Barcode technology for Cell Surface Protein protocol (#CG000186 Rev D) with the Chromium Single Cell 5’ Library & Gel Bead and Chromium Single Cell 5’ Feature Barcode Library kits (10x Genomics, #PN-1000014, #PN-1000020, #PN-1000080, #PN-1000009, #PN-1000084) according to the manufacturer’s instructions. Sequencing was performed on NovaSeq 6000 platform (Illumina).

### Quality control, normalization, and integration of scRNAseq data

The human reference used to map sequenced reads was taken from Ensembl version 102 ^107^ and prepared using 10x Cell Ranger 5.0.1 Software (*mkref* tool). The count matrices were generated by 10x Cell Ranger 5.0.1 Software (*count* tool) in either R2-only or paired-end mode. Afterwards, they were pre-processed using Seurat 4.3.1 package ^108^ on R 4.2.1. Any cell that was not marked by any expected combination of hashtags was removed. All cells with more than 10% of genes mapping to mitochondrial genes, those expressing less than 200 genes and those that were marked as doubles according to the V(D)J (more than 2 TCRα or more than 2 non-productive TCRβ or more than 1 productive TCRβ sequence present) were removed. Mitochondrial genes, ribosomal genes, genes encoding TCR variable segments (any gene symbol containing the TR[AB][VDJ] substring) and genes present in less than 3 cells were removed. Each data set was then normalized (default method and scale factor = 1 × 10^4^), scaled, subjected to dimensional reduction (PCA with 20 principal components followed by UMAP) and Louvain clustering. PTPRC counts were generated by IDEIS tool and normalized by centered-log ratio method ^28^.

Preprocessed datasets were merged, normalized (default method and scale factor = 1 × 10^4^), scaled and used for dimensional reduction (PCA with 12 principal components and UMAP) and Louvain clustering. Clusters of dead cells and any individual cells with less than 500 detected genes or more than 10% of mitochondrial genes were excluded. Integration of datasets from different batches was performed using STACAS (v 2.1.3) ^109^.

### Analysis and visualization of scRNAseq data

Transcriptional regulatory interactions were estimated using the decoupleR package (v2.10.0) using the CollecTRI database ^110^. GSEA pathways were processed using gene sets from datasets GSE9650 ^35^ and GSE11057 ^34^ with the R package fgsea (v1.20.0) ^111^. Heatmaps were created with package pheatmap (v 1.0.12). Sankey plots for figures S3 and S4 were created using RAWGraphs (v2.0) ^112^.

### HLA Typing and allele frequency analysis

We used the raw fastq files for estimation of the HLA genotype using arcasHLA ^113^. In cases where samples were profiled twice for initial and final dataset and/or for two timepoints, fastq files were merged to one fastq file per patient. These fastq files were processed with the commands *genotype* and *merge*. Reference data from the global population of the Czech Republic were obtained from The Allele Frequency Net Database ^45^ using the tool HLAfreq ^114^.

### Analysis of published data

Count matrices from the RNA sequencing data were obtained from the Gene Expression Omnibus (GEO) (GSE237218 ^37^, GSE123658 (unpublished), GSE10586 ^36^) and processed with DESeq2 ^115^. Raw fastq files from the Single Cell 3’ sequencing of PBMC of four Finnish children at risk of developing Type 1 diabetes and their gender age and HLA matched control children were obtained from the European Genome-Phenome Archive (EGAD00001005768 ^20^) and mapped with CellRanger software (10x Genomics, cellranger-7.1.0) to the *GRCh38* human reference genome. Count matrices were processed with Seurat similarly to our data. Processed scRNAseq data of PBMCs from 46 T1DM cases and 31 matched controls was obtained from Synapse under the accession code syn53641849 ^23^. Processed scRNAseq data of PBMCs from 5 T1M donors and 3 healthy donors was obtained from GEO GSE221297 ^25^. Processed scRNAseq data of PBMCs from 12 T1D donors and 12 healthy donors was obtained from ParseBio (Supplemental Table 1). Genes from all datasets were ranked by the fold changes obtained from comparing patients with T1D to healthy donors (DESeq LogFC). For comparison between datasets, the ranks were converted to 20-quantiles.

For the genotype-expression analysis of BTN3A2, we used bulk RNA sequencing data from GSE237218 ^37^, GSE123658 (unpublished), EGAD00001005767 (Kallionpaa), and scRNAseq data from the current study and from the HPAP repository. In all cases, raw fastq files were processed using using arcasHLA as described above to obtain HLA genotypes for all the donors. HLA alleles were typed to the level of two fields, which distinguishes specific HLA proteins. The information about the expression of BTN3A2 for each of the donors was obtained from the count matrices that were generated for each dataset as described above. The expression of BTN3A2 for the donors in each dataset was scaled to a range from −1 to 1 to allow comparison between datasets, i.e. the donor with the lowest expression of BTN3A2 in each dataset had a value of −1 and the donor with the highest expression in each dataset had a value of 1. The effect of study, diabetes status and particular alleles in all MHC-I and MHC-II loci was calculated using generalized linear model with gaussian distribution. The analysis was performed for each locus separately.

### Annotation of cells with previously published signatures

For annotation of cells with the TR3-56 signature, count matrix from RNA sequencing of the flow sorted CD3^+^ 56^+^ cells, NK cells, CD3^+^CD56^-^, and CD8^+^ cells was obtained from the Gene Expression Omnibus (GSE106082) ^16^. CD4 and CD8 datasets were annotated using SingleR ^116^.

For annotation of Treg cells, the differentially expressed genes from dataset GSE15659 ^48^ were retrieved from the mSigDB (#M3563). The function AddModuleScore from Seurat package was used to calculate the expression of the genes in the Treg dataset.

### Analysis of scRNAseq data from the HPAP database

From the HPAP repository, we used a collection of scRNAseq samples from splenocytes and lymph nodes of the deceased healthy or T1D donors that were processed using the HPAP CITEseq: Dual-index 3’ HT scRNAseq with Antibody Derived Tags and Hashtag Oligos protocol. The full protocol can be accessed at the HPAP database. For our analysis, raw fastq files from were downloaded from the HPAP web and mapped to the GRCh38-2020-A human reference genome using cellranger-7.1.0. Count matrices were merged and subjected to normalization (scale factor = 1 × 10^4^), scaling and dimensional reduction (PCA with 15 principal components on 2000 variable features and UMAP) and Louvain clustering. Clusters representing dead cells or contaminating cell types other than T or NK cells were removed. For analysis of the Treg subpopulations, splenic Tregs were extracted from the whole dataset and subjected to new to normalization (scale factor = 1 × 10^4^), scaling and dimensional reduction (PCA with 10 principal components on 1000 variable features and UMAP) and Louvain clustering. Batch effect of cells from different experiments was removed using STACAS (v 2.1.3).

### Analysis of TCR repertoires

TCR repertoires of the CD4^+^ and CD8^+^ T cells from initial and final experiments were profiled using 10x Chromium Single Cell V(D)J Enrichment Kit, Mouse T Cell (#PN-1000071) and mapped by the 10x Cellranger 5.0.1 sotware (*vdj* tool) to human reference obtained the International ImMunoGeneTics Information System (IMGT) ^117^ for the immune receptor repertoire profiling. Additional V(D)J sequences were extracted from the gene expression library using the MiXCR 3.0.12 software ^118^. For the clonal expansion analyses, cells with the same nucleotide sequences of the CDR3α and CDR3β were considered clones. In cases where we detected two productive rearrangements of CDR3α, the cells were considered the same clones if the nucleotide sequences of CDR3β and both CDR3α were shared, or if the nucleotide sequences of CDR3β and one CDR3α were shared while the second CDR3α was not detected.

For the repertoire overlap analysis, unique amino-acid sequences of the CDR3α or CDR3β were extracted for each donor. Then, the percentage of overlapping sequences in one participant with all of the other participants was calculated and quantified in the following comparisons: i) healthy donors with other healthy donors, ii) healthy donor with a T1D donor, or iii) two T1D donors. In addition, the Self T1 – Self T0 comparisons were calculated as the overlap between CDR3α or CDR3β sequences in the same donor in the two timepoints.

For the analysis of the length of the CDR3β, unique amino-acid sequences of the CDR3β were extracted for each donor. The lengths of the sequences were quantified as the frequency of each length in a particular donor and visualized in T1D patients and healthy donors.

For analysis of the biochemical properties of the CDR3 sequences, we used the R package Peptides ^119^. For each of the biochemical properties available in the package, the values for all the unique CDR3β sequences were calculated and compared between healthy donors and T1D patients. Each sequence was taken into the analysis the number of times it occurred in the dataset, i.e. a CDR3β sequence representing an expanded clone that was detected in three donors with 50 occurrences in donor 1, 10 occurrences in donor 2 and 1 occurrence in donor 3 would be counted 61 times. To see the global biases of the repertoires in T1D patients and healthy donors not influenced by the clonal expansion, we calculated the biochemical properties for all the unique CDR3β sequences in each donor, and averaged these values per donor.

For all the analyses except for clonal expansion, the analysis was performed on conventional cells only, i.e., MAIT cells, NK cells, and γδT T cells were excluded from the CD8^+^ dataset, and iNKT cells were excluded from the CD4^+^ dataset.

The analyses of CDR3β length and biochemical properties were performed also on samples from splenocytes of the non-diabetic or T1D donors from the HPAP dataset. In the HPAP repository, the TCR libraries were prepared as described previously ^120^. Briefly, T cell receptor beta chain gene rearrangements were bulk sequenced from gDNA using custom primers. Typically 100 ng of input DNA was amplified per replicate. Sequencing libraries were prepared using 2 × 300 bp paired-end kits (Illumina MiSeq Reagent Kit v3, 600-cycle, Illumina Inc., San Diego, CA). The raw sequencing data (fastq files) were downloaded from the HPAP database and mapped to the using Mixcr v4.5.0 to the human reference obtained from International ImMunoGeneTics Information System (IMGT) ^117^.

### Antibodies for flow cytometry

The following antibodies were used for flow cytometry staining: anti-CD3 (clone MEM-57, Exbio #A7-202-T100), anti-CD3 (clone UCHT1, BD #561007), anti-CD4 (clone SK3, Biolegend #344632), anti-CD4 (clone OKT4, Biolegend #317433), anti-CD16 (clone B73.1, Biolegend #360729), anti-CD8 (clone MEM-31, Exbio #1A-207-T100), anti-CD25 (clone M-A251, Biolegend #356146), anti-CD45RA (clone MEM-56, Exbio #1F-223-T100), anti-CD45RO (clone UCHL1, Biolegend #304232), anti-CD45R/B220 (clone RA3-6B2, Biolegend #103248), anti-CD56 APC-R700 (clone NCAM16.2, BD #565140), anti-CD127 (clone A019D5, Biolegend #351333), anti-CD137 (clone 4B4, eBioscience #25-1379-42), anti-CD183 (clone G025H7, Biolegend #353705), anti-CD184 (clone 12G5, Exbio #1P-146-T100), anti-CD196 (clone G034E3, Biolegend #353429), anti-CD197 (clone G043H7, Biolegend #353227), anti-CD226 (clone 11A8, Exbio #1P-926-T100), anti-EOMES (clone X4-83, BD Pharmingen #567167), anti-FOXP3 (clone 206D, Biolegend #320125), anti-IκBα (clone L35A5, Cell Signaling #5743S), anti-Granzyme B (clone QA16A02, Biolegend #372213), anti-Granzyme K (clone GM26E7, Biolegend #370513), anti-HLA-DR (clone LN3, Biolegend #327019), anti-Ki-67 (clone Ki-67, Biolegend #350535), anti-TCR gamma/delta (clone 11F2, Exbio #T7-912-T100), anti-TCR Vα24-Jα18 (clone 6B11, Biolegend #342929).

### Flow cytometry analysis

Flow cytometry was performed in two batches. In each batch, aliquoted PBMCs stored in the liquid nitrogen were gently thawed by slow, sequential dilution in RPMI medium containing 10% FBS. Cells were counted and the same amount of cells (1.2 million in batch one, 2 million in batch two) were used for staining. The counted cells were divided into three equal parts. The first part was used for extracellular staining of live cells, the other two parts were used for intracellular staining of fixed cells. To prevent unspecific binding to Fc receptors, Human TrueStain FcX (BioLegend #422301) was added to all staining mixes. For staining of extracellular markers (Panel 1: CD4 OKT4 BV421, CD16 BV510, TCR Vα24-Jα18 BV605, CD45RO BV650, CD197 BV711, CD45RA FITC, CD3 UCHT1 PE-Cy5, HLA-DR PerCP/Cy5.5, CD183 PE, CD25 PE/Fire700, CD196 PE/Dazzle 594, TCR gamma/delta PE-Cy7, CD8 APC, CD56 APC-R700), cells were incubated with diluted antibodies and LIVE/DEAD Fixable Near-IR Dead Cell Stain Kit (Invitrogen #L34975) for 30 min on ice immediately after isolation. For staining of intracellular markers and transcription factors, cells were first stained for 30 min on ice with the mix containing LIVE/DEAD Fixable Near-IR Dead Cell Stain Kit (Invitrogen #L34975) and extracellular markers (CD3 MEM-57 AF700, CD4 SK3 BV421, CD8 APC, CD25 PE/Fire700, CD45RA FITC, CD45RO BV650, CD45R/B220 BV510) and then fixed and permeabilized using the eBioscience Foxp3 / Transcription Factor Staining Buffer Set (Invitrogen #00-5523-00) according to the manufacturer’s instructions, washed, and stained overnight with antibodies for intracellular markers in 4°C (Panel 2: extracellular markers and CD127 BV605, Ki-67 BV750, HLA-DR PerCP/Cy5.5, CD226 PE, FOXP3 PE/Dazzle 594, CD137 Pe/Cy7; Panel 3: extracellular marekrs and CD127 BV605, Ki-67 BV750, IκBα AF488, GZMK PerCP/Cy5.5, CD184 PE, Eomes PE/CF594, GZMB PE/Cy7).

Flow cytometry was carried out with Cytek Aurora flow cytometer, configuration 4L 16V-14B-10YG-8R (Cytek). Data were analyzed using FlowJo software v10.10 (BD Life Sciences). Anomalies in flow rate, signal acquisition, and dynamic range were removed using FlowJo plugin FlowAI ^121^.

### HPAP flow cytometry

Flow cytometry data (fcs files) from PBMC of patients with T1D and healthy donors were downloaded from the HPAP. These samples were collected from deceased donors on the date of death and analyzed either fresh or after cryopreservation. For our analysis, we used only PBMC samples from patients with T1D and healthy donors. Two samples were excluded due to suspected duplicity and six samples were excluded based on the lack of compensation metadata in the fcs file. Samples were analyzed in groups based on three different panels for staining and manually compensated when needed. We analyzed 41 samples in Panel 1 – CD4 phenotyping panel, 19 samples in Panel 2 – CD8 phenotyping panel and 26 samples in Panel 3 – CD8 phenotyping panel focused primarily on antigen-specific cells. In cases when the same patient had measurements in multiple panels, these values were averaged. For analysis, we excluded patients youger than 5 years as appropriate age-matched heatlhy controls were not available.

### Statistical analysis

The statistical analysis was performed using tests indicated in the Figure legends using R v4.2.1. Quantitative scRNAseq and flow cytometry data were tested using the nonparametric Mann–Whitney test without a correction for multiple comparisons or, for a comparison of the same patients at two different timepoints, using paired Wilcoxon signed rank test without a correction for multiple comparisons. Two-tailed tests were performed. The number of biological replicates (cells and donors) is indicated in the respective Figure legends.

### Bayesian statistics

Differences in population composition were assessed by a Bayesian generalized linear model with a negative binomial response, with an offset term to normalize for the total number of cells in a sample. The input data consisted the counts of cells for each Level 3 population and the model included a fixed effect for the type of sample (healthy, T1D at time 1, T1D at time 0) and the Level 3 population. Additionally, random effects of the type of sample were added, grouped by both Level 2 and Level 3 populations. The shape parameter was also predicted, with a fixed effect for Level 2 populations and a random intercept for Level 3 populations. The model was fitted with the brms package ^122^.

## Data availability

Processed scRNAseq data are deposited on Zenodo, DOI: 10.5281/zenodo.14222417. Raw sequencing data are not deposited to protect the privacy of the donors.

## Code availability

The scripts used for data analysis are found here: https://github.com/Lab-of-Adaptive-Immunity/dia.

## Acknowledgement

We gratefully acknowledge Ladislav Cupak for technical assistance, Zdenek Cimburek and Matyas Sima (flow cytometry facility, IMG) for cell sorting, Sarka Kocourkova and Michal Kolar (Laboratory of Genomics and Bioinformatics, IMG) for cDNA library preparations, and Stepanka Pruhova, Stanislava Kolouskova, Barbora Obermannova, Lenka Drnkova and Lukas Plachy who cared for the children with T1D during the course of the study, and Jachym Antonin Harwood for advice on the manuscript. VNi and BC are students of the Faculty of Science, Charles University in Prague.

This project was supported by the Czech Science Foundation (22-21356S to OS), project National Institute of Virology and Bacteriology (Programme EXCELES, LX22NPO5103 to OS)—funded by the European Union—Next Generation EU, Charles University Grant Agency (252208 to VN), European Union’s Horizon 2020 research and innovation programme under grant agreement No. 802878 (ERC Starting Grant FunDiT to OS), core funding provided by the Institute of Molecular Genetics of the Czech Academy of Sciences (RVO 68378050), and core funding provided by the Institute of Molecular Genetics of the Czech Academy of Sciences (RVO 68378050), and Czech Ministry of Health (conceptual support project to research organization 00064203 – FN Motol). Computational resources were provided by the e-INFRA CZ project (ID:90254), supported by the Ministry of Education, Youth and Sports of the Czech Republic. This manuscript used data acquired from the Human Pancreas Analysis Program (HPAP-RRID:SCR_016202) Database (https://hpap.pmacs.upenn.edu/), a Human Islet Research Network (RRID:SCR_014393) consortium (UC4-DK-112217, U01-DK-123594, UC4-DK-112232, and U01-DK-123716).

## Author contribution

VNe and ZS enrolled human donors and provided biological samples and metadata, AN processed and cryopreserved the biological samples and prepared cells for scRNAseq experiments, VNi and JM analyzed transcriptomic and TCR profiling data generated in this study, VNi analyzed publicly available transcriptomic data, VNi and BC performed and analyzed flow cytometry experiments, BC analyzed HPAP flow cytometry data, MM performed Bayesian statistical analysis, ZS and OS supervised the project and provided administrative tasks required for the study, VNi and OS wrote the manuscript. All authors revised the manuscript.

