## Supplemental for "An imbalance of naïve and effector T-cell phenotypes in early type 1 diabetes across conventional and regulatory subsets"

**Table S1.** List of transcriptomics datasets from public resources and previously published studies used for validation of the findings from the current cohort. [1-5]

| | Study | Reposited data | Participants | Cell type | Method |
|---|---|---|---|---|---|
| Bulk transcriptomics | Newman et al., 2023 [3] | GSE237218 | CD4+/CD25+ T cells: 49 T1D cases, 35 controls<br>CD4+/CD25– T cells: 53 T1D cases, 52 controls<br>Memory CD4+ T cells: 19 T1D cases, 27 controls | Sorted T cell subsets | Bulk RNAseq |
| | Valentim et al., 2018 * | GSE123658 | 39 T1D cases, 43 healthy donors | Whole blood, hemoglobin depleted | Bulk RNAseq |
| | Kallionpää et al., 2019 [2] | EGAD00001005767 | 7 T1D cases, 8 controls | Sorted T cell subsets | Bulk RNAseq |
| | Jailwala et al., 2009 [1] | GSE10586 | 12 T1D cases, 15 healthy donors | Sorted T cell subsets | Microarray |
| Single-cell transcriptomics | HPAP - scRNAseq * | https://hpap.pmacs.upenn.edu/ | 4 T1D cases, 8 controls | Splenocytes | scRNAseq |
| | Kallionpää et al., 2019 [2] | EGAD00001005768 | 4 T1D cases, 4 controls | PBMC | scRNAseq |
| | Honardoost et al., 2024 [4] | syn53641849 | 46 T1D cases, 31 controls | PBMC | scRNAseq |
| | ParseBio * | https://resources.parsebiosciences.com/dataset-wt-mega-one-million-pbmc-type-1-diabetes | 12 T1D, 12 healthy donors | PBMC | scRNAseq |
| | Zhong et al., 2024 [5] | GSE221297 | 5 new onset, 3 healthy controls | PBMC | scRNAseq |
| TCR profiling | HPAP - TCRseq * | https://hpap.pmacs.upenn.edu/ | 16 T1D cases, 38 controls | Splenocytes | TCRseq |
| FACS | HPAP - FACS * | https://hpap.pmacs.upenn.edu/ | 23 T1D cases, 32 controls | PBMC | FACS |

* Unpublished dataset

**Figure S1.**

Peripheral blood from fasting blood sampling was analyzed by routine clinical testing.

A) Levels of glycosylated hemoglobin (left) and C-peptide (right) in fasting blood samples of healthy donors, children with T1D at diagnosis and the same patients at one-year follow-up. P-value was calculated using the Kruskal-Wallis test. Bar at median. n = 11 Healthy, n = 29 T1D T0, n = 28 T1D T1.
B) Correlation of the levels of fasting C-peptide from blood samples of T1D patients at T0 and the levels of C-peptide sampled at a random timepoint (left), or with the levels of fasting C-peptide sampled from the same patients at T1. P-value was calculated using the Pearson's correlation test. n = 28 T1D donors.
C) Percentages (upper panels) and absolute counts (lower panels) of major white blood cell populations of healthy donors and T1D at T0 and T1. P-value was calculated using the Kruskal-Wallis test. Bar at median. n = 11 Healthy, n = 29 T1D T0, n = 28 T1D T1.

Figure S1

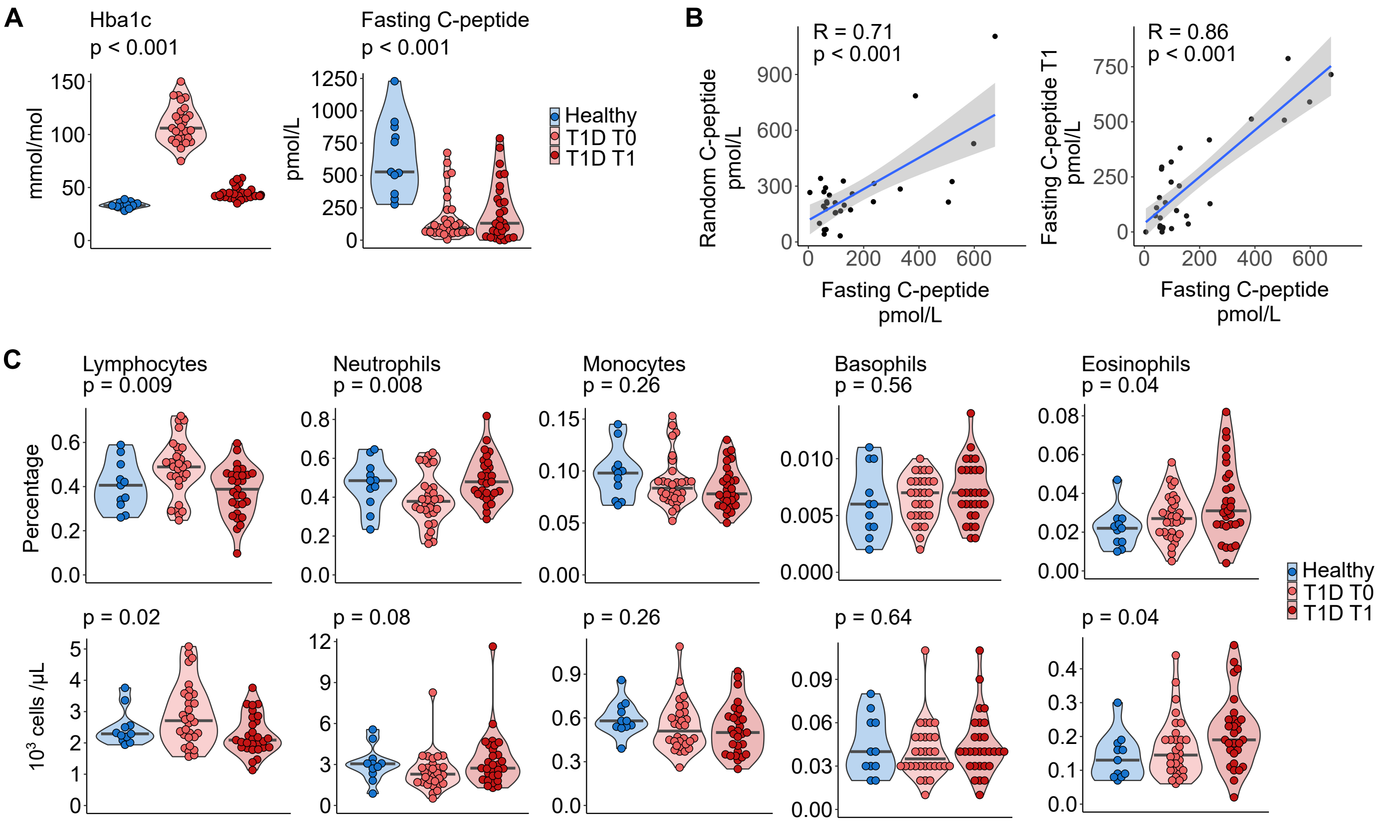
A
Hba1c
p < 0.001
mmol/mol
Fasting C-peptide
p < 0.001
pmol/L
Healthy
T1D T0
T1D T1
B
R = 0.71
p < 0.001
Random C-peptide pmol/L
Fasting C-peptide pmol/L
R = 0.86
p < 0.001
Fasting C-peptide T1 pmol/L
C
Lymphocytes
p = 0.009
Neutrophils
p = 0.008
Monocytes
p = 0.26
Basophils
p = 0.56
Eosinophils
p = 0.04
Percentage
p = 0.02
p = 0.08
p = 0.26
p = 0.64
p = 0.04
10^3 cells /μL

**Figure S2.**
(A-D) An initial experiment in which $CD4^+$ and $CD8^+$ T cells were sorted from 12 T1D children and 6 healthy donors.

(A) Representative gating used for sorting of $CD4^+$ and $CD8^+$ T cells from fresh frozen PBMCs.
(B) UMAP plot showing $CD8^+$ T cells from initial experiments before quality control and removal of contaminating cells. Clusters of cells were annotated as naïve (blue) or antigen-experienced (AgExp, red) based on marker genes and isoforms of CD45 (PTPRC) inferred from gene expression data. n = 18,270 cells from 18 donors.
(C) UMAP plot showing $CD4^+$ T cells from initial experiments before quality control and removal of contaminating cells. Clusters of cells were divided into naïve (blue) and AgExp (red) based on marker genes and isoforms of CD45 (PTPRC) inferred from gene expression data. n = 14278 cells from 18 donors
(D) Quantification of naïve and AgExp clusters in the initial $CD4^+$ and $CD8^+$ T-cell datasets.

(E-H) Final experiment in which AgExp enriched $CD4^+$ and $CD8^+$ T cells were sorted from 30 T1D donors and 13 healthy donors. For each patient, naïve and AgExp cells were sorted at 1:5 ratio.

(E) Representative gating used for sorting of AgExp enriched $CD4^+$ and $CD8^+$ T cells from fresh frozen PBMC.

(F) The UMAP plot shows $CD8^+$ T cells from final experiments before quality control and removal of contaminating cells. Clusters of cells were annotated as naïve (blue) or AgExp (red) based on marker genes and isoforms of PTPRC inferred from gene expression data. n = 93,412 cells from 43 donors
(G) The UMAP plot shows $CD4^+$ T cells from the final experiment before the quality control and removal of contaminating cells. Clusters of cells were divided into naïve (blue) and AgExp (red) based on marker genes and isoforms of PTPRC inferred from gene expression data. n = 78,635 cells from 43 donors
(H) Quantification of naïve and AgExp clusters in the final $CD4^+$ and $CD8^+$ T-cell datasets.

(I) The quality control and filtering of cells from the initial and final experiments in the $CD8^+$ T-cell dataset (top) and $CD4^+$ T-cell dataset (bottom). From left are shown: the UMAP plots of all cells before quality control with Louvain clusters; DotPlots showing the expression of canonical T-cell genes and genes characteristic for contaminating cell types in the clusters; Violin plots showing percentages of mitochondrial genes and total gene counts in all clusters together with filtering criteria (red = removed, white = kept); and UMAP plots showing the cells that passed the quality control (grey) or were removed (red).

(J) The PCA analysis showing the batch effect correction using STACAS in the initial and final experiments after the quality control. Each dot represents a sample from one donor. Colors represent different batches.

Figure S2

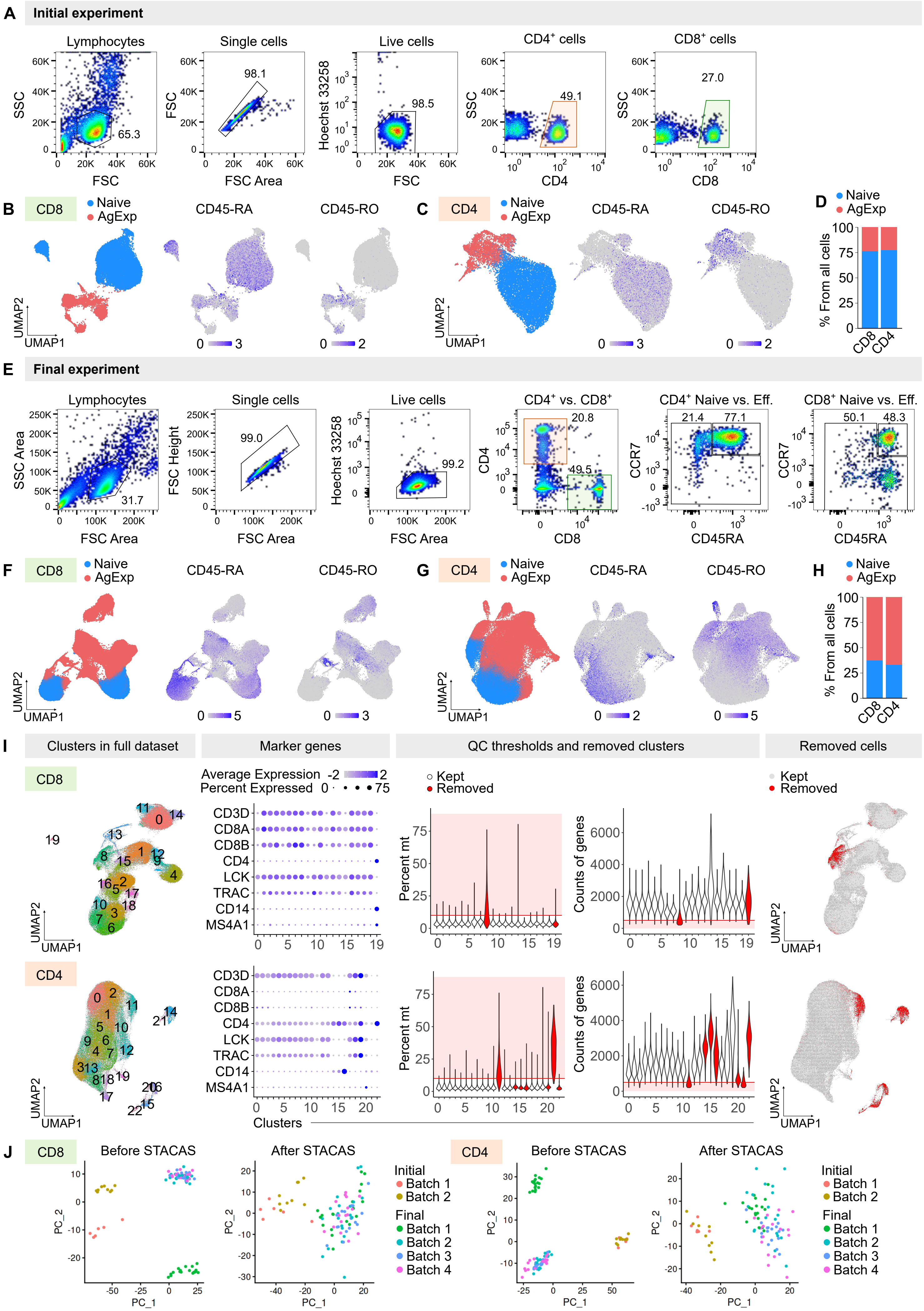
A
Initial experiment
Lymphocytes
Single cells
Live cells
CD4+ cells
CD8+ cells
SSC
FSC
Hoechst 33258
CD4
CD8
65.3
98.1
98.5
49.1
27.0
B
CD8
Naive
AgExp
CD45-RA
CD45-RO
UMAP1
UMAP2
C
CD4
D
% From all cells
E
Final experiment
CD4+ vs. CD8+
CD4+ Naive vs. Eff.
CD8+ Naive vs. Eff.
SSC Area
FSC Area
FSC Height
CCR7
CD45RA
31.7
99.0
99.2
20.8
49.5
21.4
77.1
50.1
48.3
F
G
H
I
Clusters in full dataset
Marker genes
QC thresholds and removed clusters
Removed cells
Average Expression
Percent Expressed
Kept
Removed
CD3D
CD8A
CD8B
CD4
LCK
TRAC
CD14
MS4A1
Percent mt
Counts of genes
Clusters
J
Before STACAS
After STACAS
Initial
Batch 1
Batch 2
Final
Batch 3
Batch 4
PC_1
PC_2

**Figure S3.**

A) The UMAP projection of $CD8^+$ T cells showing the unconventional subsets and NK cells, which were not included in the main analysis shown in Fig. 1C-E. Louvain clusters were merged based on the functional relevance. n = 95,229 cells from 43 donors.

B) The same UMAP projection as in (A) showing the detection of TCR CDR3α sequences typical for MAIT cells [6] in the top panels and the expression of TRDC and TRGV9 gene segments in the bottom panels.

C) The heatmap shows the relative expression of the marker genes that characterize clusters presented in (A). For comparison, clusters of conventional $CD8^+$ T cells (Fig. 1C) were also included. Colors represent row-scaled z-score of average expression of a gene in a cluster.

D) The heatmap shows the transcriptional regulation of clusters presented in (A). The transcription factors were identified as the key regulators based on differential expression in each cluster using the CollecTRI transcriptional regulons database [7]. For comparison, clusters of conventional $CD8^+$ T cells (Fig. 1C) were also included.

E) The heatmap shows the transcriptional regulation of clusters presented in Fig. 1C. The transcription factors were identified as the key regulators based on differential expression in each cluster using CollecTRI transcriptional regulons database [7].

F) The Sankey plot shows the $CD8^+$ T-cell populations (Level 1) and their subclustering into the main clusters (Level 2) and subclusters (Level 3). Sizes of the dots represent the relative abundance of the cluster.

Figure S3

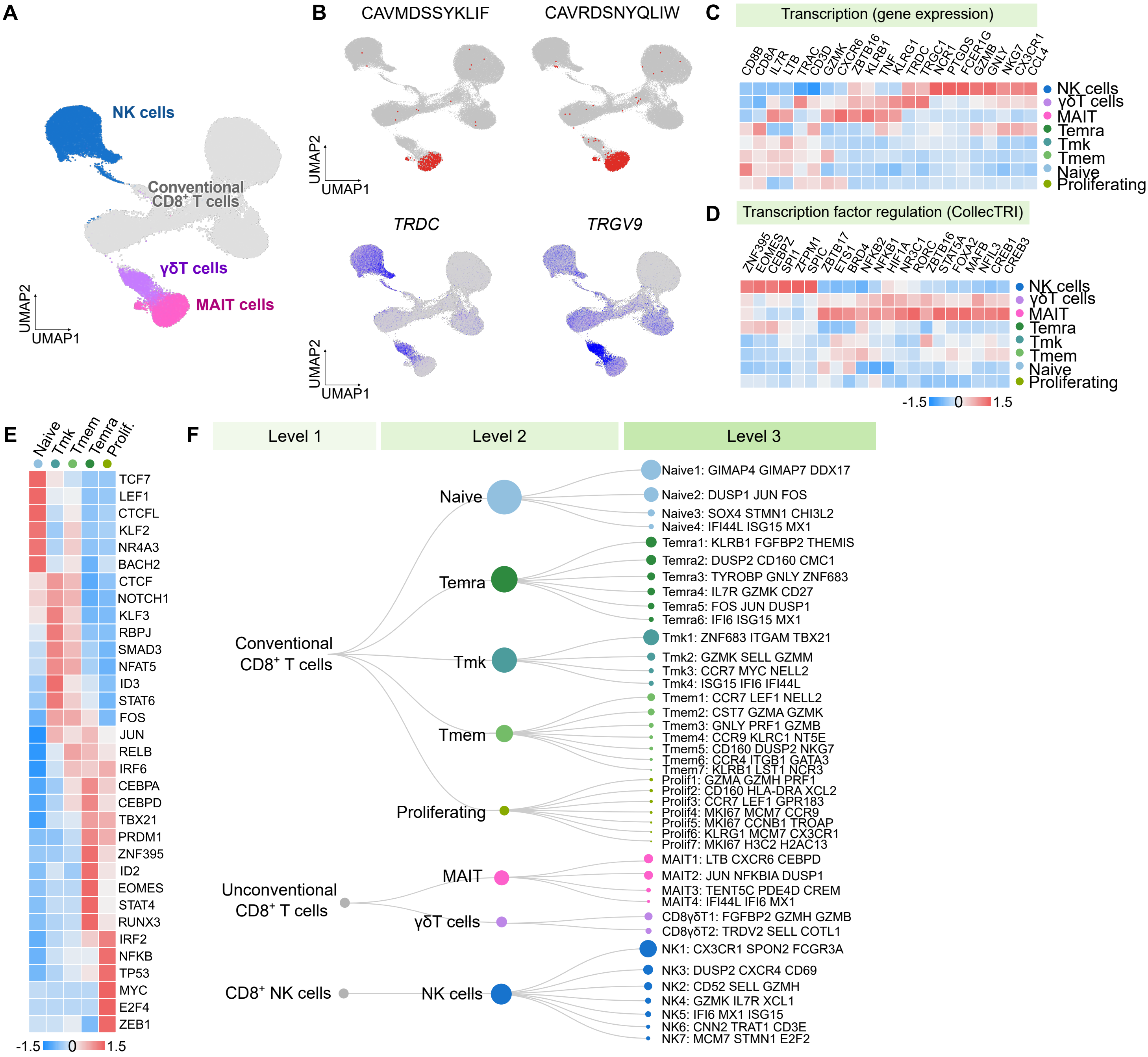

A
NK cells
Conventional CD8+ T cells
γδT cells
MAIT cells
UMAP2
UMAP1
B
CAVMDSSYKLIF
CAVRDSNYQLIW
TRDC
TRGV9
C
Transcription (gene expression)
CD8B CD8A IL7R LTB TRAC CD3D GZMK CXCR6 ZBTB16 KLRB1 TNF KLRG1 TRDC TRGC1 NCR1 PTGDS FCER1G GZMB GNLY NKG7 CX3CR1 CCL4
NK cells
γδT cells
MAIT
Temra
Tmk
Tmem
Naive
Proliferating
D
Transcription factor regulation (CollecTRI)
ZNF395 EOMES CEBPZ SP11 ZFPM1 SPIC ZBTB17 ETS1 BRD4 NFKB2 NFKB1 HIF1A NR3C1 RORC ZBTB16 STAT5A FOXA2 MAFB NFIL3 CREB1 CREB3
-1.5 0 1.5
E
Naive Tmk Tmem Temra Prolif.
TCF7 LEF1 CTCFL KLF2 NR4A3 BACH2 CTCF NOTCH1 KLF3 RBPJ SMAD3 NFAT5 ID3 STAT6 FOS JUN RELB IRF6 CEBPA CEBPD TBX21 PRDM1 ZNF395 ID2 EOMES STAT4 RUNX3 IRF2 NFKB TP53 MYC E2F4 ZEB1
F
Level 1
Level 2
Level 3
Conventional CD8+ T cells
Unconventional CD8+ T cells
CD8+ NK cells
Naive
Temra
Tmk
Tmem
Proliferating
MAIT
γδT cells
NK cells
Naive1: GIMAP4 GIMAP7 DDX17
Naive2: DUSP1 JUN FOS
Naive3: SOX4 STMN1 CHI3L2
Naive4: IFI44L ISG15 MX1
Temra1: KLRB1 FGFBP2 THEMIS
Temra2: DUSP2 CD160 CMC1
Temra3: TYROBP GNLY ZNF683
Temra4: IL7R GZMK CD27
Temra5: FOS JUN DUSP1
Temra6: IFI6 ISG15 MX1
Tmk1: ZNF683 ITGAM TBX21
Tmk2: GZMK SELL GZMM
Tmk3: CCR7 MYC NELL2
Tmk4: ISG15 IFI6 IFI44L
Tmem1: CCR7 LEF1 NELL2
Tmem2: CST7 GZMA GZMK
Tmem3: GNLY PRF1 GZMB
Tmem4: CCR9 KLRC1 NT5E
Tmem5: CD160 DUSP2 NKG7
Tmem6: CCR4 ITGB1 GATA3
Tmem7: KLRB1 LST1 NCR3
Prolif1: GZMA GZMH PRF1
Prolif2: CD160 HLA-DRA XCL2
Prolif3: CCR7 LEF1 GPR183
Prolif4: MKI67 MCM7 CCR9
Prolif5: MKI67 CCNB1 TROAP
Prolif6: KLRG1 MCM7 CX3CR1
Prolif7: MKI67 H3C2 H2AC13
MAIT1: LTB CXCR6 CEBPD
MAIT2: JUN NFKBIA DUSP1
MAIT3: TENT5C PDE4D CREM
MAIT4: IFI44L IFI6 MX1
CD8γδT1: FGFBP2 GZMH GZMB
CD8γδT2: TRDV2 SELL COTL1
NK1: CX3CR1 SPON2 FCGR3A
NK3: DUSP2 CXCR4 CD69
NK2: CD52 SELL GZMH
NK4: GZMK IL7R XCL1
NK5: IFI6 MX1 ISG15
NK6: CNN2 TRAT1 CD3E
NK7: MCM7 STMN1 E2F2

**Figure S4.**

A) The UMAP projection of $CD4^+$ T cells showing the unconventional cells, which were not included in the main analysis shown in Fig. 1F-H. Louvain clusters were merged based on functional relevance. n = 79,876 cells from 43 donors.

B) The same UMAP projection as in (A) showing the detection of TRA gene segments TRAV10 recombined with TRAJ18, which is typical for iNKT cells (left), and the CDR3α sequence typical for iNKT cells [6] (middle). The expression of *ZBTB16* is shown (right).

C) The heatmap shows the relative expression of marker genes that characterize clusters presented in (A). For comparison, clusters of conventional $CD4^+$ T cells (Fig. 1F) were also included. Colors represent row-scaled z-score of average expression of a gene in a cluster.

D) The heatmap shows the transcriptional regulation of the unconventional cells in comparison to conventional clusters. Transcription factors were identified as the key regulators based on differential expression in each cluster using the CollecTRI transcriptional regulons database [7].

E) Identification of reads mapping specifically to CD45-RA (PTPRC isoform RA) by the IDEIS software [8].

F) The heatmap shows the transcriptional regulation of clusters presented in Fig. 1F. The transcription factors were identified as the key regulators based on differential expression in each cluster using the CollecTRI transcriptional regulons database [7].

G) The Sankey plot shows the $CD4^+$ populations (Level 1) and their subclustering into the main clusters (Level 2) and subclusters (Level 3). Sizes of the dots represent relative abundance of the cluster.

Figure S4

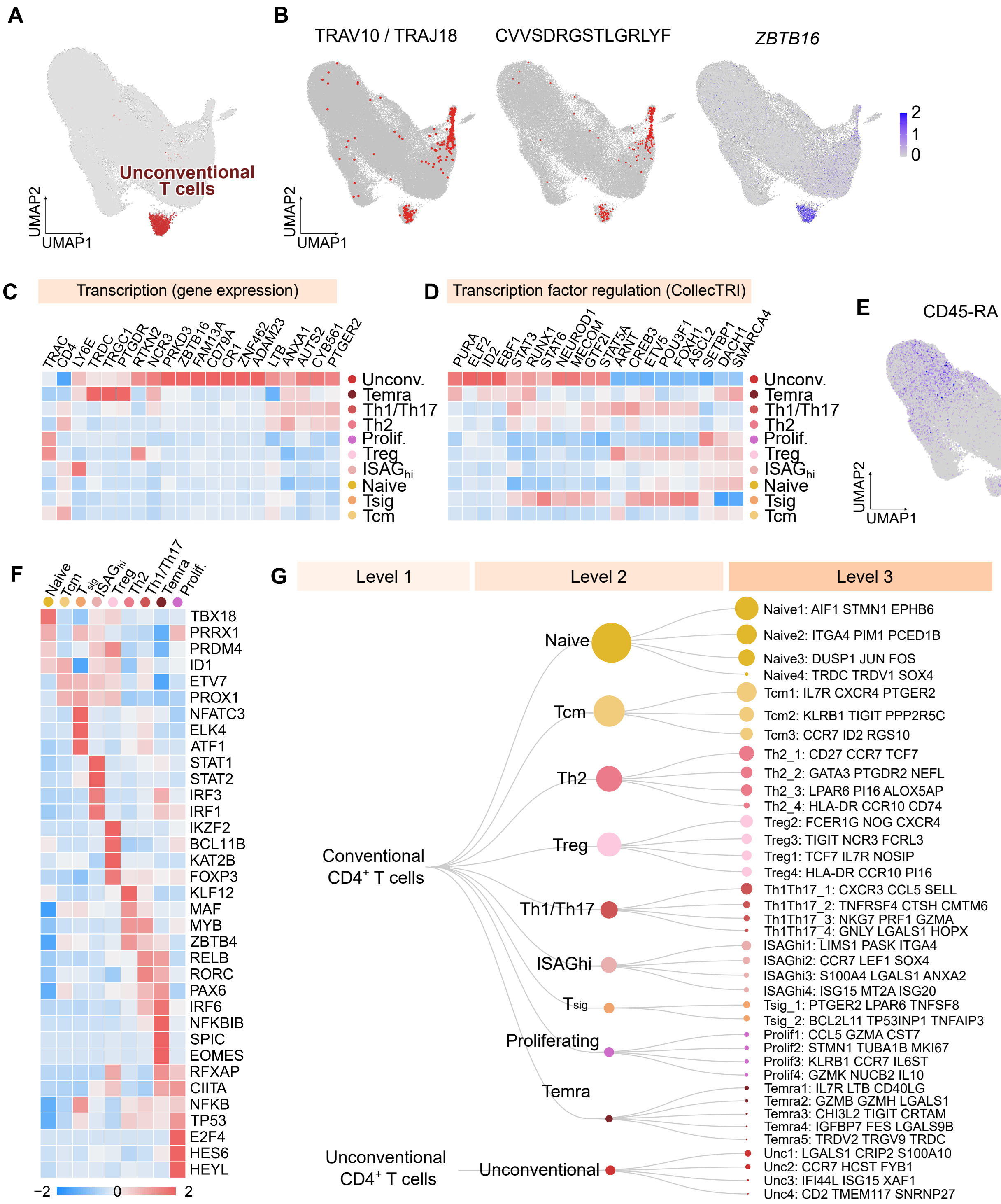

A
Unconventional T cells
UMAP2
UMAP1
B
TRAV10 / TRAJ18
CVVSDRGSTLGRLYF
ZBTB16
2
1
0
C
Transcription (gene expression)
TRAC CD4 LY6E TRDC TRGC1 PTGDR RTKN2 NCR3 PRKD3 ZBTB16 FAM13A CD79A CR1 ZNF462 ADAM23 LTB ANXA1 AUTS2 CYB561 PTGER2
Unconv.
Temra
Th1/Th17
Th2
Prolif.
Treg
ISAGhi
Naive
Tsig
Tcm
D
Transcription factor regulation (CollecTRI)
PURA ELF2 ID2 EBF1 STAT3 RUNX1 STAT6 NEUROD1 MECOM GTF2I STAT5A ARNT CREB3 ETV5 POU3F1 FOXH1 ASCL2 SETBP1 DACH1 SMARCA4
E
CD45-RA
5
0
F
Naive Tcm Tsig ISAGhi Treg Th2 Th1/Th17 Temra Prolif.
TBX18
PRRX1
PRDM4
ID1
ETV7
PROX1
NFATC3
ELK4
ATF1
STAT1
STAT2
IRF3
IRF1
IKZF2
BCL11B
KAT2B
FOXP3
KLF12
MAF
MYB
ZBTB4
RELB
RORC
PAX6
IRF6
NFKBIB
SPIC
EOMES
RFXAP
CIITA
NFKB
TP53
E2F4
HES6
HEYL
−2
0
2
G
Level 1
Level 2
Level 3
Conventional CD4+ T cells
Unconventional CD4+ T cells
Naive
Tcm
Th2
Treg
Th1/Th17
ISAGhi
Tsig
Proliferating
Temra
Unconventional
Naive1: AIF1 STMN1 EPHB6
Naive2: ITGA4 PIM1 PCED1B
Naive3: DUSP1 JUN FOS
Naive4: TRDC TRDV1 SOX4
Tcm1: IL7R CXCR4 PTGER2
Tcm2: KLRB1 TIGIT PPP2R5C
Tcm3: CCR7 ID2 RGS10
Th2_1: CD27 CCR7 TCF7
Th2_2: GATA3 PTGDR2 NEFL
Th2_3: LPAR6 PI16 ALOX5AP
Th2_4: HLA-DR CCR10 CD74
Treg2: FCER1G NOG CXCR4
Treg3: TIGIT NCR3 FCRL3
Treg1: TCF7 IL7R NOSIP
Treg4: HLA-DR CCR10 PI16
Th1Th17_1: CXCR3 CCL5 SELL
Th1Th17_2: TNFRSF4 CTSH CMTM6
Th1Th17_3: NKG7 PRF1 GZMA
Th1Th17_4: GNLY LGALS1 HOPX
ISAGhi1: LIMS1 PASK ITGA4
ISAGhi2: CCR7 LEF1 SOX4
ISAGhi3: S100A4 LGALS1 ANXA2
ISAGhi4: ISG15 MT2A ISG20
Tsig_1: PTGER2 LPAR6 TNFSF8
Tsig_2: BCL2L11 TP53INP1 TNFAIP3
Prolif1: CCL5 GZMA CST7
Prolif2: STMN1 TUBA1B MKI67
Prolif3: KLRB1 CCR7 IL6ST
Prolif4: GZMK NUCB2 IL10
Temra1: IL7R LTB CD40LG
Temra2: GZMB GZMH LGALS1
Temra3: CHI3L2 TIGIT CRTAM
Temra4: IGFBP7 FES LGALS9B
Temra5: TRDV2 TRGV9 TRDC
Unc1: LGALS1 CRIP2 S100A10
Unc2: CCR7 HCST FYB1
Unc3: IFI44L ISG15 XAF1
Unc4: CD2 TMEM117 SNRNP27

**Figure S5.**

A) Quantification of the cluster composition for clusters shown in Fig. 1C. The violin plots show the percentage of cells in each subcluster from total conventional $CD8^+$ T cells in healthy and T1D donors at T0 and T1.

B) Quantification of the cluster composition for clusters shown in Fig. S3A. The violin plots show the percentage of cells in each subcluster from total $CD8^+$ T cells in healthy and T1D donors at T0 and T1.

C) The quantification of the cluster composition for clusters shown in Fig. 1F. The violin plots show the percentage of cells in each subcluster from total conventional $CD4^+$ T cells in healthy and T1D donors at T0 and T1.

D) The quantification of the unconventional $CD4^+$ cells shown in Fig. S4A. Violin plots show the percentage of unconventional $CD4^+$ T cells from total $CD4^+$ T cells in healthy and T1D donors at T0 and T1.

E) The Bayesian analysis of the abundance of subsets of $CD8^+$ T cells in T1D donors at T0 vs. healthy donors (left), T1D donors at T1 vs. T0 (middle) and T1D donors at T1 vs. healthy donors.

F) The Bayesian analysis of the abundance of subsets of $CD4^+$ T cells in T1D donors at T0 vs. healthy donors (left), T1D donors at T1 vs. T0 (middle) and T1D donors at T1 vs. healthy donors.

G) Correlation of the levels of fasting C-peptide from T1D donors at T1 with the frequency of their populations of $CD8^+$ T cells at T0.

H) Correlation of the levels of fasting C-peptide from T1D donors at T1 with the frequency of their populations of $CD4^+$ T cells at T0.

A-D - P-value was calculated using two-tailed Mann-Whitney test between healthy donors and T1D T0, or two-tailed paired Mann-Whitney test between T1D donors at T0 and T1. Bar at median.

Figure S5

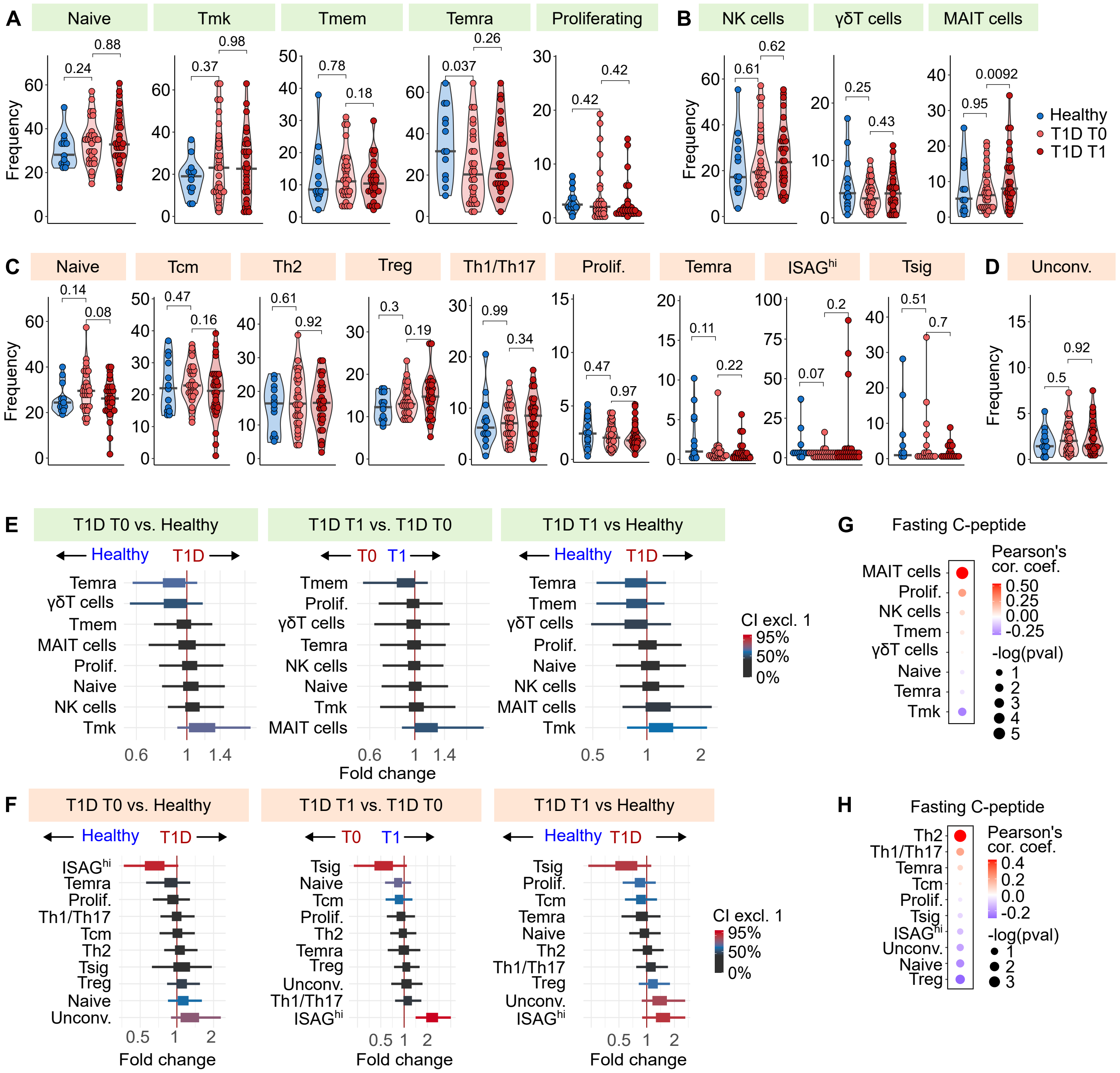

A
Naive
Tmk
Tmem
Temra
Proliferating
B
NK cells
γδT cells
MAIT cells
Healthy
T1D T0
T1D T1
Frequency
C
Naive
Tcm
Th2
Treg
Th1/Th17
Prolif.
Temra
ISAGhi
Tsig
D
Unconv.
E
T1D T0 vs. Healthy
T1D T1 vs. T1D T0
T1D T1 vs Healthy
Healthy
T1D
T0
T1
CI excl. 1
95%
50%
0%
Fold change
F
T1D T0 vs. Healthy
T1D T1 vs. T1D T0
T1D T1 vs Healthy
Fold change
G
Fasting C-peptide
Pearson's cor. coef.
-log(pval)
MAIT cells
Prolif.
NK cells
Tmem
γδT cells
Naive
Temra
Tmk
H
Fasting C-peptide
Pearson's cor. coef.
-log(pval)
Th2
Th1/Th17
Temra
Tcm
Prolif.
Tsig
ISAGhi
Unconv.
Naive
Treg

**Figure S6.**

A) The volcano plot shows differentially expressed genes in $CD4^+$ T cells (orange) and $CD8^+$ T cells (green) from T1D donors at T0 to healthy donors (left) or T1D donors at T1 to healthy donors (right). The average log2 fold changes were calculated using the FindMarkers function from the Seurat package (Wilcoxon test) using the Wilcoxon test and default criteria. Multiple-hypothesis testing is controlled using the Bonferroni correction.

B) Differentially expressed genes between healthy donors and T1D donors at T0 and T1. The average log2 fold changes were calculated using the FindMarkers function from the Seurat package (Wilcoxon test). Genes with adjusted p-value < 0.05 were plotted and selected genes were manually labelled for $CD4^+$ T cells and their subsets as shown in Fig. 1F (top) and for $CD8^+$ T cellsand their subsets as shown in Fig. 1C (bottom). Multiple-hypothesis testing is controlled using the Bonferroni correction.

C) The volcano plot shows differentially expressed genes in $CD4^+$ T cells (orange) and $CD8^+$ T cells (green) from donors with T1D (T0) who presented with ketoacidosis at diagnosis compared to donors who did not suffer from it (left) or from donors with T1D (T0) who did not present with partial remission of the disease at one-year follow-up compared to those who did (right). The average log2 fold changes were calculated using the FindMarkers function from the Seurat package (Wilcoxon test). Multiple-hypothesis testing is controlled using the Bonferroni correction.

D) A dotplot of the difference in expression of selected genes representing markers of naïve T cells, effector T cells, Type-I interferon-responsive genes and signaling regulators in the following comparisons: T1D T0 vs healthy, T1D T1 vs healthy, T1D T0 vs T1D T1, T1D T0 without partial remission at T1 vs T1D T0 with partial remission at T1, and T1D T0 with ketoacidosis vs. T1D T0 without ketoacidosis. The average log2 fold changes were calculated in all the stated subpopulations using the FindMarkers function from the Seurat package using the Wilcoxon test and adjusted criteria: logfc.threshold = -Inf, min.pct = -Inf, min.diff.pct = -Inf, min.cells.feature = 1, return.thresh = 1. Color represents the average value of log2 fold change, size of dots represents the –log10 of the adjusted p-value. Multiple-hypothesis testing is controlled using the Bonferroni correction.

E) A dotplot showing the enrichment of selected gene sets in the following comparisons: T1D T0 vs healthy, T1D T1 vs healthy, T1D T0 vs T1D T1, T1D T0 without partial remission at T1 vs T1D T0 with partial remission at T1, and T1D T0 with ketoacidosis vs. T1D T0 without ketoacidosis. The ranks of genes per each comparison were obtained using the FindMarkers function from the Seurat package as described in (D). Color represents the value of normalized enrichment score (NES), size of dots represents the –log10 of the adjusted p-value. Multiple-hypothesis testing is controlled using the Bonferroni correction.

Figure S6

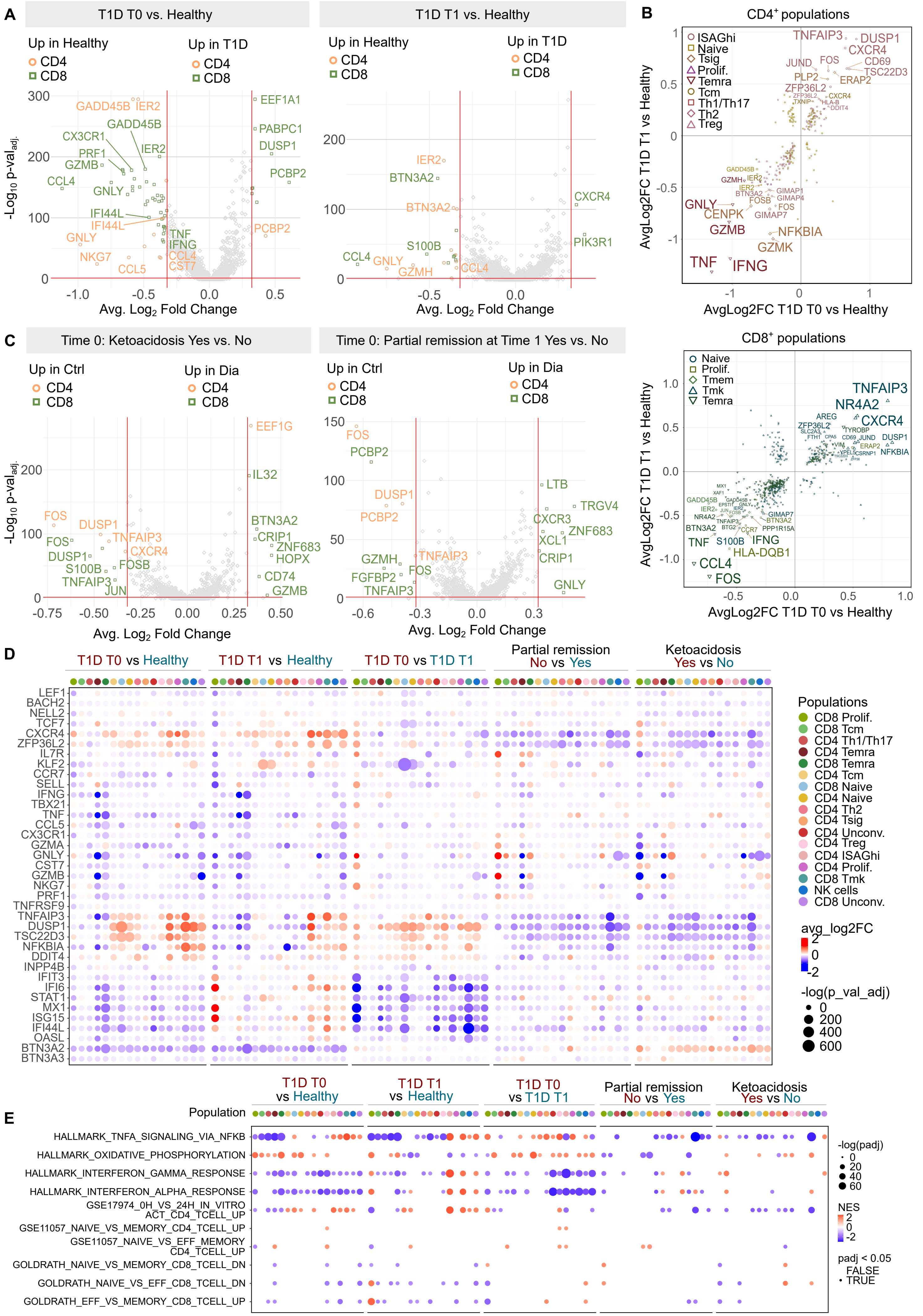
A
T1D T0 vs. Healthy
T1D T1 vs. Healthy
Up in Healthy
Up in T1D
CD4
CD8
-Log10 p-val adj.
Avg. Log2 Fold Change
B
CD4+ populations
CD8+ populations
AvgLog2FC T1D T1 vs Healthy
AvgLog2FC T1D T0 vs Healthy
C
Time 0: Ketoacidosis Yes vs. No
Time 0: Partial remission at Time 1 Yes vs. No
Up in Ctrl
Up in Dia
D
T1D T0 vs Healthy
T1D T1 vs Healthy
T1D T0 vs T1D T1
Partial remission No vs Yes
Ketoacidosis Yes vs No
Populations
avg_log2FC
-log(p_val_adj)
E
Population
HALLMARK_TNFA_SIGNALING_VIA_NFKB
HALLMARK_OXIDATIVE_PHOSPHORYLATION
HALLMARK_INTERFERON_GAMMA_RESPONSE
HALLMARK_INTERFERON_ALPHA_RESPONSE
GSE17974_0H_VS_24H_IN_VITRO ACT_CD4_TCELL_UP
GSE11057_NAIVE_VS_MEMORY_CD4_TCELL_UP
GSE11057_NAIVE_VS_EFF_MEMORY CD4_TCELL_UP
GOLDRATH_NAIVE_VS_MEMORY_CD8_TCELL_DN
GOLDRATH_NAIVE_VS_EFF_CD8_TCELL_DN
GOLDRATH_EFF_VS_MEMORY_CD8_TCELL_UP
-log(padj)
NES
padj < 0.05
FALSE
TRUE

**Figure S7.**

A) Representative gating of CD4$^+$ and CD8$^+$ Naïve and Effector cells in flow cytometry samples from the HPAP database. In the three rows, we present the same populations gated in three different panels obtained from the original source: Panel 1 – CD4 phenotyping panel, Panel 2 – CD8 phenotyping panel and Panel 3 – CD8 phenotyping panel focused on antigen-specific cells. Samples are representatives of n = 41 sample in Panel 1, n = 19 samples in Panel 2, n = 26 samples in Panel 3.

B) The heatmap shows DE genes consistently up- or down-regulated in T1D donors compared to healthy donors across different published studies. Count matrices containing transcriptomics data of T1D patients and healthy donors from different studies (see Supplemental Table 1 for details) were processed using the same pipeline to obtain fold changes for all genes in T1D patients vs healthy donors. These fold changes were used to calculate 20 quantiles for each dataset, i.e. genes in quantile 1 were the most upregulated in T1D vs healthy and genes in quantile 20 were the most upregulated in healthy. Genes which were shared among the most datasets in the quantiles 1 (left) or 20 (right) are shown.

Figure S7

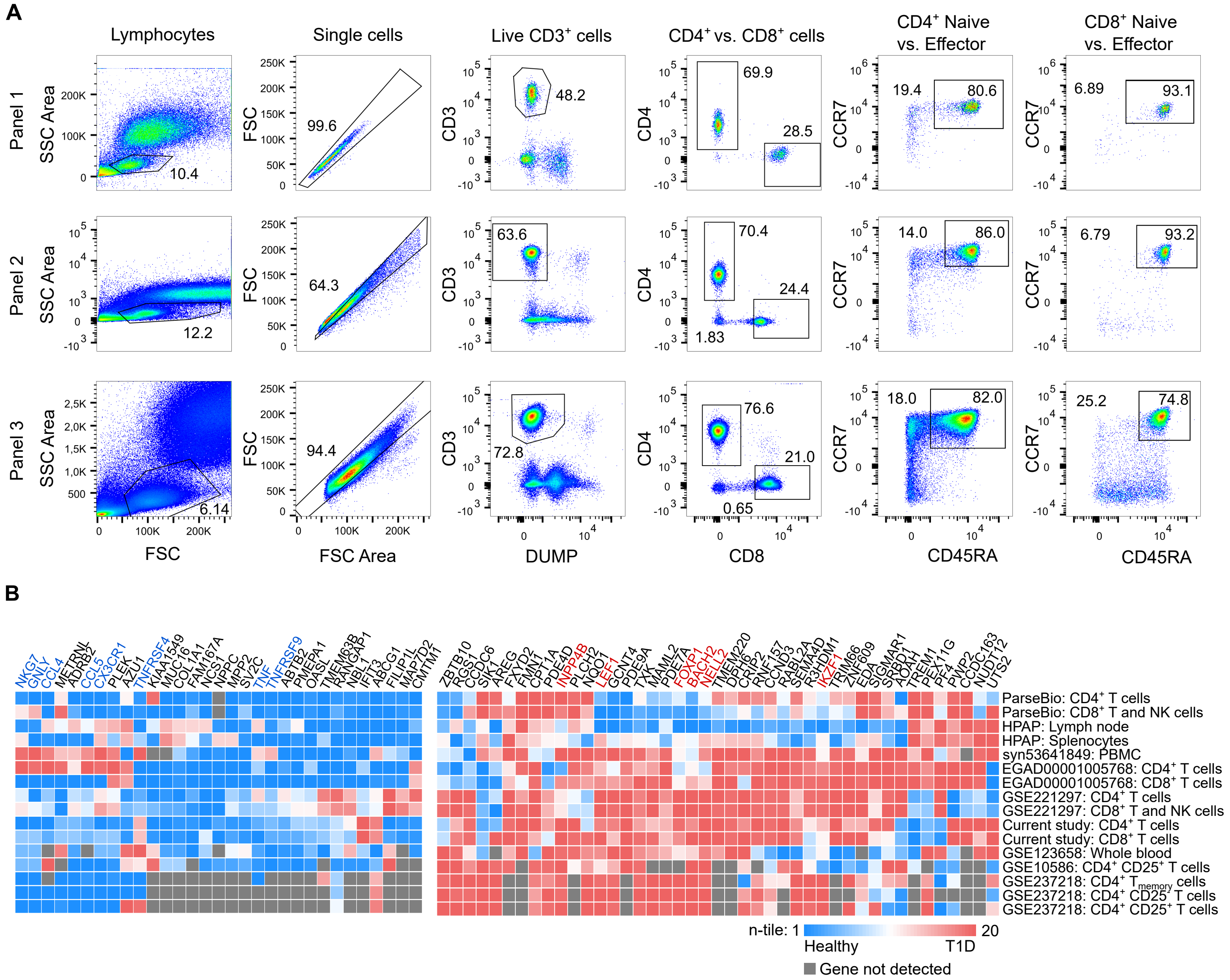
A
Lymphocytes
Single cells
Live CD3+ cells
CD4+ vs. CD8+ cells
CD4+ Naive vs. Effector
CD8+ Naive vs. Effector
Panel 1
Panel 2
Panel 3
SSC Area
FSC
CD3
CD4
CCR7
FSC Area
DUMP
CD8
CD45RA
10.4
99.6
48.2
69.9
28.5
19.4
80.6
6.89
93.1
12.2
64.3
63.6
70.4
24.4
1.83
14.0
86.0
6.79
93.2
6.14
94.4
72.8
76.6
21.0
0.65
18.0
82.0
25.2
74.8
B
ParseBio: CD4+ T cells
ParseBio: CD8+ T and NK cells
HPAP: Lymph node
HPAP: Splenocytes
syn53641849: PBMC
EGAD00001005768: CD4+ T cells
EGAD00001005768: CD8+ T cells
GSE221297: CD4+ T cells
GSE221297: CD8+ T and NK cells
Current study: CD4+ T cells
Current study: CD8+ T cells
GSE123658: Whole blood
GSE10586: CD4+ CD25+ T cells
GSE237218: CD4+ Tmemory cells
GSE237218: CD4+ CD25- T cells
GSE237218: CD4+ CD25+ T cells
n-tile: 1
20
Healthy
T1D
Gene not detected

**Figure S8.**

A) Analysis of the frequencies of HLA class I (left) and HLA class II (right) genotypes of healthy donors in the Czech Republic and participants of the current study. HLA genotypes of T1D and healthy donors were inferred from scRNAseq data using the tool ArcasHLA [9]. The frequencies of HLA alleles in the global population of the Czech Republic was obtained from The Allele Frequency Net Database. P-value was calculated using binomial test and adjusted for multiple comparison using the Bonferroni method. For testing, the frequency of the allele in global Czech population was calculated as the average of frequencies in the four Czech Republic reference datasets.

(B-C) Normalized expression of BTN3A2 was compared among five studies with T1D and healthy donors. HLA genotype of donors was obtained from the original metadata or inferred from the raw RNAseq data using ArcasHLA. The effect of study, diabetes status, and particular alleles in all MHC-I and MHC-II loci was calculated using generalized linear model with gaussian distribution. Statistically significant predictor variables in the model are marked in red color. The analysis was performed for each locus separately.

B) The effect of disease status and each of the listed studies on BTN3A2 expression.
C) The effect of the MHC-I and MHC-II alleles on BTN3A2 expression.

Figure S8

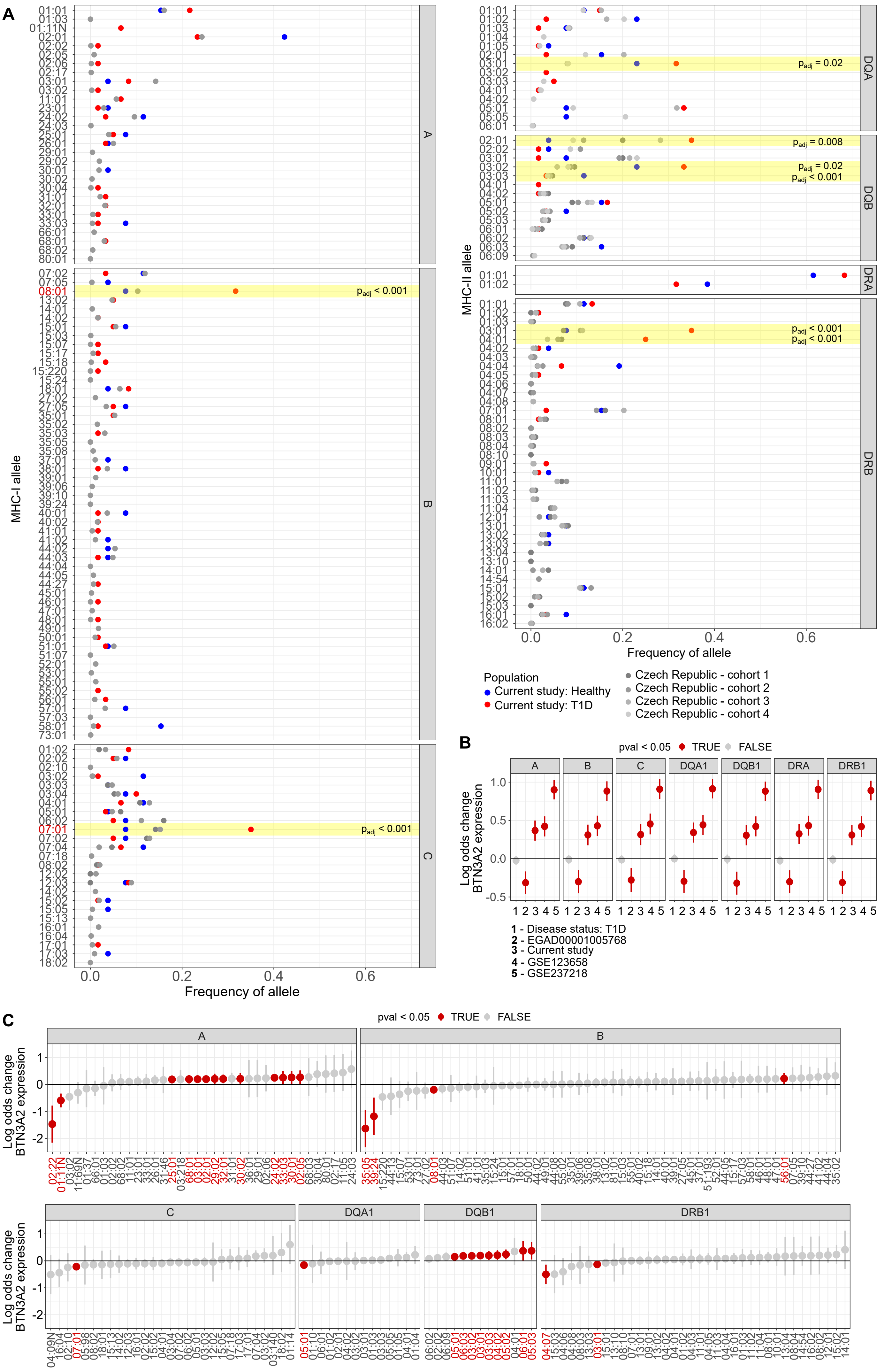
A
MHC-I allele
MHC-II allele
Frequency of allele
$p_{adj}$ < 0.001
$p_{adj}$ = 0.02
$p_{adj}$ = 0.008
DQA
DQB
DRA
DRB
Population
Current study: Healthy
Current study: T1D
Czech Republic - cohort 1
Czech Republic - cohort 2
Czech Republic - cohort 3
Czech Republic - cohort 4
B
pval < 0.05 TRUE FALSE
A
B
C
DQA1
DQB1
DRA
DRB1
Log odds change BTN3A2 expression
1 - Disease status: T1D
2 - EGAD00001005768
3 - Current study
4 - GSE123658
5 - GSE237218
C

**Figure S9.**

A) Quantification of the cluster composition for clusters shown in Fig 3A. Violin plots show the percentage of cells in each subcluster from total Treg cells in healthy and T1D donors at T0 and T1. Bar at median. P-value was calculated using two-tailed Mann-Whitney test between healthy and T1D T0 donors, or two-tailed paired Mann-Whitney test between T1D donors at T0 and T1.

B) Correlation of the levels of fasting C-peptide measured at T1 and the frequency of Treg1 (left) or Treg4 (right) subsets at T0. P-value was calculated using the Pearson's correlation test. n = 28 T1D donors.

(C-E) Analysis of Treg population extracted from scRNAseq data from published datasets: GSE221297 (top), ParseBio (middle) and HPAP (bottom). Samples in GSE221297 were obtained from PBMC of 5 T1D patients and 3 healthy donors. Samples in ParseBio were obtained from PBMC of 12 T1D patients and 12 healthy donors. Samples in HPAP were obtained from splenocytes of 4 deceased donors with T1D and 8 without T1D.

(C) UMAP projection of the Treg cells from a particular dataset. Cells are colored by Louvain clusters.

(D) A density plot showing the difference between the distribution of Treg cells in samples coming from healthy and T1D donors.

(E) The heatmap of marker genes that characterize clusters presented in (C). The color represents row-scaled z-score of average expression of a gene in a cluster.

(F) A quantification of the percentage of cells in each cluster (clusters are shown in C) in each sample. In the boxplots, the line represents median, the hinges correspond to the first and third quartiles, the whiskers represent 1.5x inter-quartile range.

(G) A quantification of the average percentage of cells in each Treg sub-cluster in healthy donors (blue) and T1D donors (red) for each dataset. Concerning our study, the clusters Treg1-4 were matched to clusters TregA-D here. P-value was calculated using the paired t-test.

Figure S9

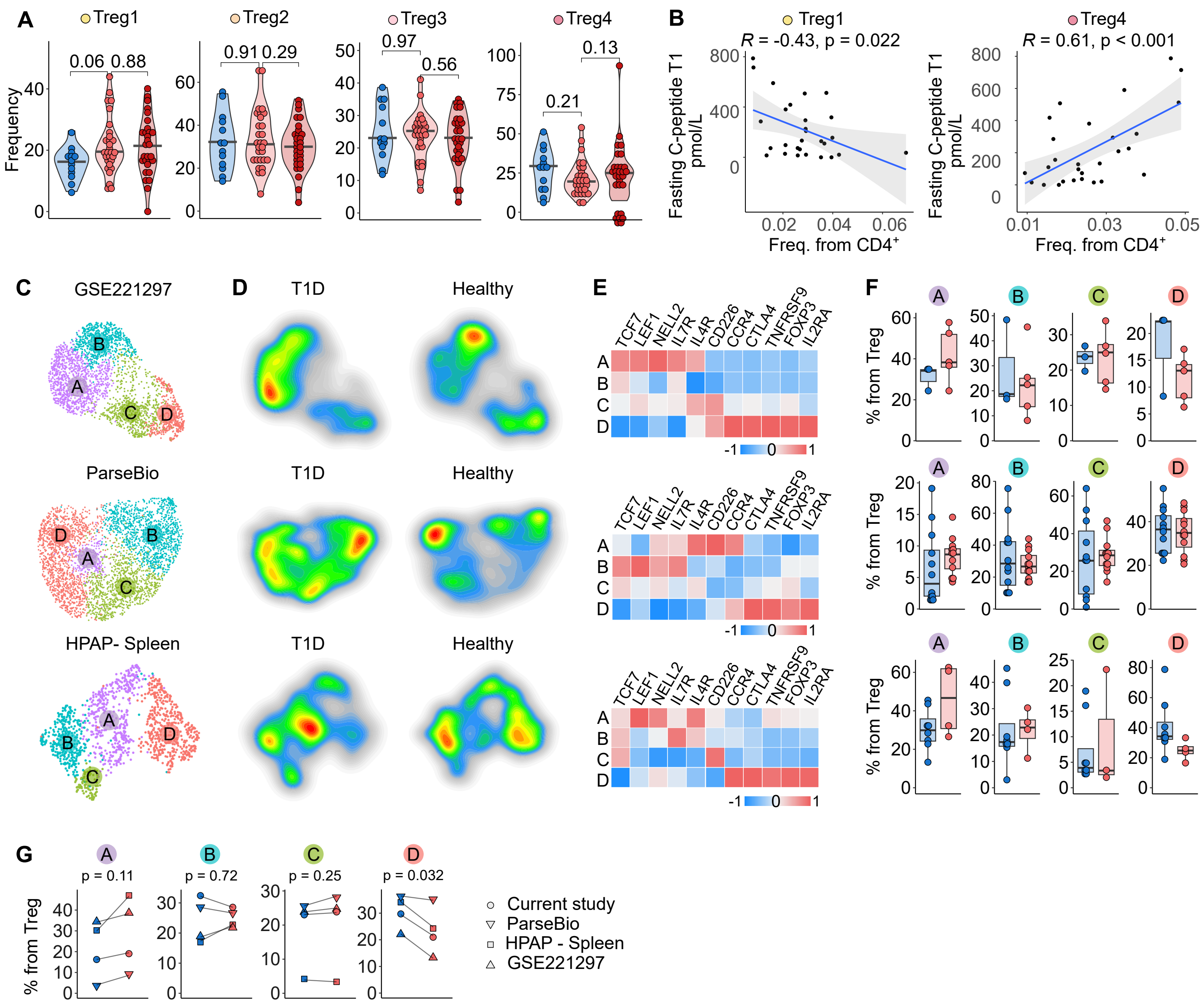
A
Treg1
Treg2
Treg3
Treg4
Frequency
0.06 0.88
0.91 0.29
0.97
0.56
0.21
0.13
B
Treg1
R = -0.43, p = 0.022
Treg4
R = 0.61, p < 0.001
Fasting C-peptide T1 pmol/L
Freq. from CD4+
C
GSE221297
ParseBio
HPAP- Spleen
D
T1D
Healthy
E
TCF7
LEF1
NELL2
IL7R
IL4R
CD226
CCR4
CTLA4
TNFRSF9
FOXP3
IL2RA
F
% from Treg
G
p = 0.11
p = 0.72
p = 0.25
p = 0.032
% from Treg
Current study
ParseBio
HPAP - Spleen
GSE221297

**Figure S10.**
(A-B) Flow cytometry analysis of the Treg cells and Treg subpopulations in the current study. n = 13 healthy donors, n = 27 T1D T0 donors, n = 28 T1D T1 donors.

A) Representative flow cytometry gating of the Treg cells and Treg subpopulations in the current study.
B) Quantification of the Treg cells and Treg subpopulations in donors with T1D and healthy donors measured by flow cytometry. The following populations are quantified: Foxp3$^+$ from CD4$^+$ T cells, CD45RA$^+$ from Foxp3$^+$ CD4$^+$ T cells, 4-1BB$^+$ CD45RO$^+$ from Foxp3$^+$ CD4$^+$ T cells, CD45RA$^+$ CD226$^+$ CD127$^+$ from Foxp3$^+$ CD4$^+$ T cells. Representative gating is shown in (A). P-value was calculated using two-tailed Mann-Whitney test between healthy and T1D T0 donors, or two-tailed paired Mann-Whitney test between T1D donors at T0 and T1.

(C-D) Flow cytometry analysis of the Treg cells and Treg subpopulations in the HPAP dataset. n = 21 non-diabetic donors, n = 17 T1D donors.

C) Representative flow cytometry gating of the Treg cells and Treg subpopulations in the HPAP dataset.
D) Quantification of the Foxp3$^+$ from CD4$^+$ T cells and CD45RA$^+$ Foxp3$^+$ CD4$^+$ T cells in donors with T1D and healthy donors from the HPAP database. Representative gating is shown in (C). P-value was calculated using two-tailed Mann-Whitney test. Bar at median.

Figure S10

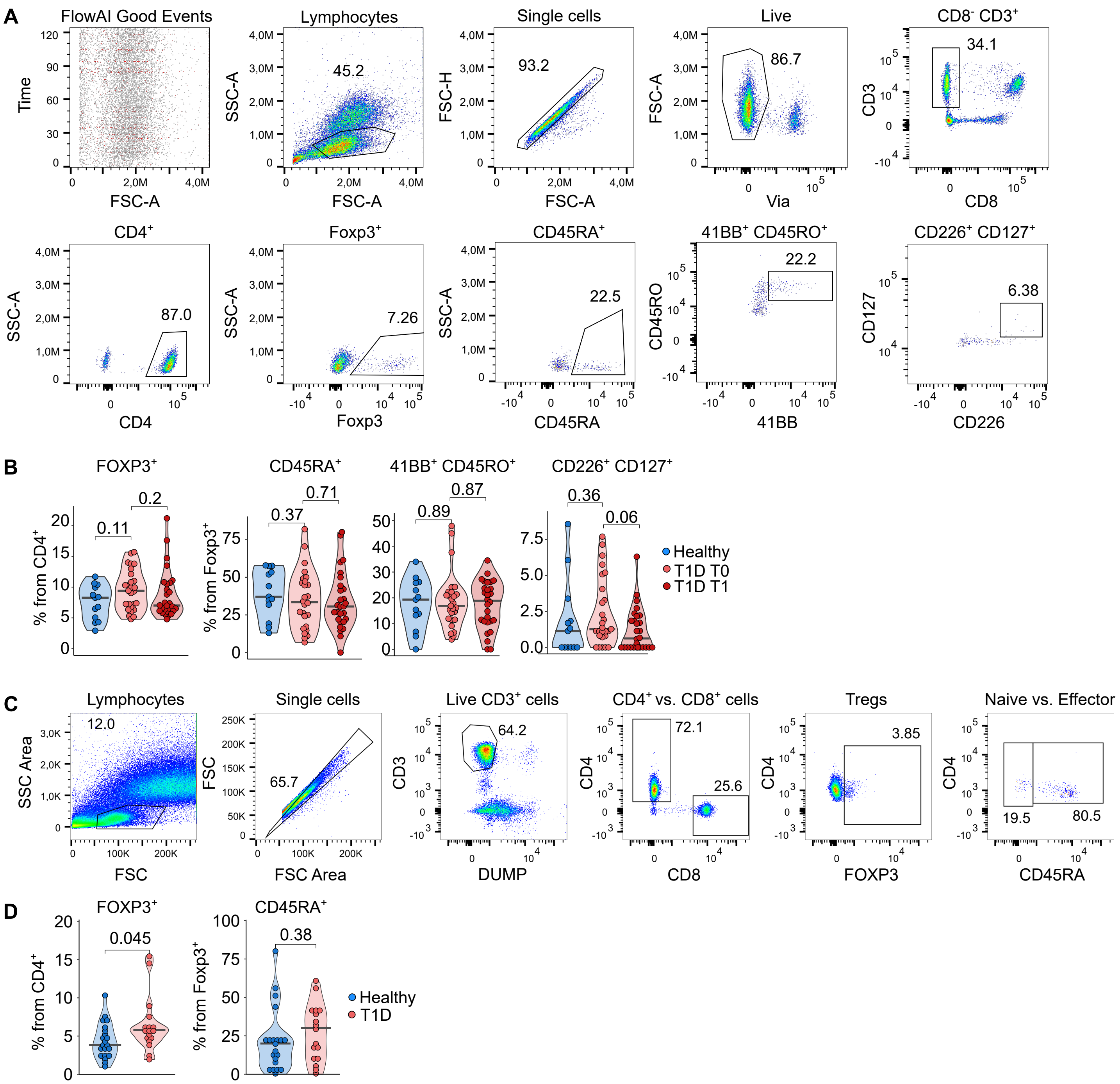
A
FlowAI Good Events
Time
FSC-A
Lymphocytes
SSC-A
45.2
Single cells
FSC-H
93.2
Live
Via
86.7
CD8- CD3+
CD3
CD8
34.1
CD4+
CD4
87.0
Foxp3+
Foxp3
7.26
CD45RA+
CD45RA
22.5
41BB+ CD45RO+
CD45RO
41BB
22.2
CD226+ CD127+
CD127
CD226
6.38
B
FOXP3+
% from CD4+
0.11
0.2
CD45RA+
% from Foxp3+
0.37
0.71
41BB+ CD45RO+
0.89
0.87
CD226+ CD127+
0.36
0.06
Healthy
T1D T0
T1D T1
C
Lymphocytes
SSC Area
FSC
12.0
Single cells
FSC Area
65.7
Live CD3+ cells
CD3
DUMP
64.2
CD4+ vs. CD8+ cells
CD4
CD8
72.1
25.6
Tregs
FOXP3
3.85
Naive vs. Effector
CD45RA
19.5
80.5
D
FOXP3+
% from CD4+
0.045
CD45RA+
% from Foxp3+
0.38
Healthy
T1D

**Figure S11.**

A) Cells from the datasets GSE221297 and ParseBio were annotated with a previously published dataset of FACS-sorted TR3-56 ($CD3^+$ $CD56^-$) cells, NK cells, $CD3^+$ CD56-, and $CD8^+$ cells (GSE106082 [10] using the SingleR package. The violin plots show quantification of the annotation scores of TR3-56 cells (i.e., similarity of the gene expression of a particular cell to that of TR3-56 cells) in $CD8^+$ and NK or $CD4^+$ T cells in healthy and T1D donors. GSE221297 $CD4^+$: n = 35,050 cells from 8 donors, GSE221297 $CD8^+$ and NK: n = 38,243 cells from 8 donors, ParseBio $CD4^+$: n = 63,219 cells from 24 donors, ParseBio $CD8^+$ and NK: n = 49,844 cells from 24 donors. P-value was calculated using two-tailed Mann-Whitney test. Bar at median.

B) UMAP projection of $CD8^+$ T cells showing groups of cells used for the analysis of unconventional $CD8^+$ subsets. Louvain clusters were merged based on functional relevance. n = 95,229 cells from 43 donors.

C) Violin plots showing the percentage of cells from the clusters indicated in (B) that have non-zero expression of the selected gene within the specified cluster. Data is based on expression profiles from 87 samples. Bar at median.

D) Percentage of cells in each cluster indicated in (B) annotated as TR3-56 cells.

E) Volcano plot showing differentially expressed genes in Unconventional $CD8^+$ T cells from patients with T1D sampled at their diagnosis compared to healthy donors (left) or from one-year follow-up samples of the same patients compared to healthy donors (right). The average log2 fold changes were calculated using the FindMarkers function from the Seurat package (Wilcoxon test). Multiple-hypothesis testing is controlled using the Bonferroni correction.

F) Representative flow cytometry gating of the populations of cells shown in Fig. 4G-H.

G) Flow cytometry analysis of $CD8^{low}$ T cells from the PBMC of deceased donors with T1D compared to healthy donors from the HPAP database. Representative flow cytometry gating of $CD8^{low}$ T cells in the HPAP dataset.

H) Quantification of $CD8^{low}$ T cells in T1D patients and healthy donors. P-value was calculated using two-tailed Mann-Whitney test.

Figure S11

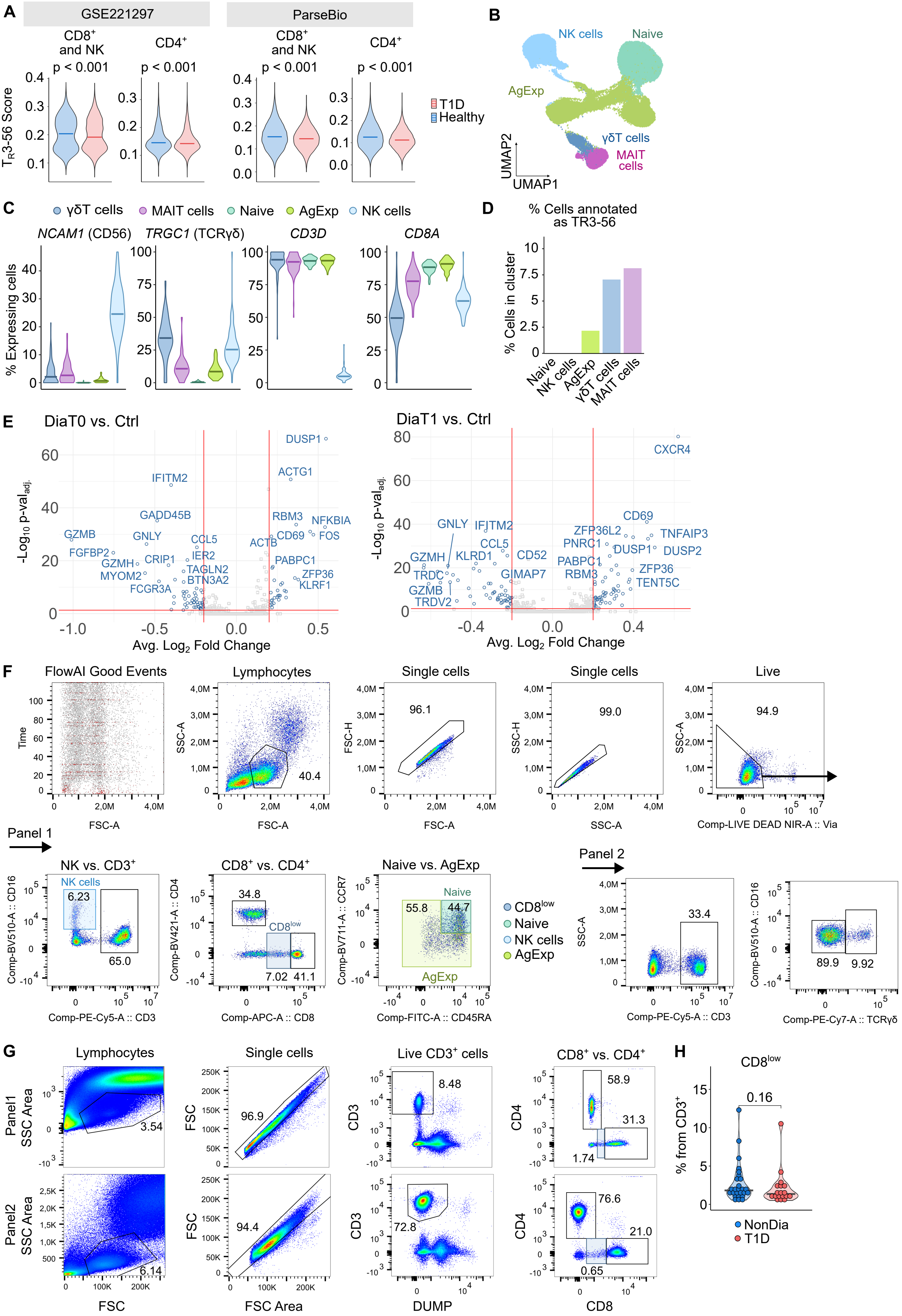

A
GSE221297
ParseBio
CD8+ and NK
CD4+
p < 0.001
T1D
Healthy
TR3-56 Score
B
NK cells
Naive
AgExp
γδT cells
MAIT cells
UMAP1
UMAP2
C
γδT cells
MAIT cells
Naive
AgExp
NK cells
NCAM1 (CD56)
TRGC1 (TCRγδ)
CD3D
CD8A
% Expressing cells
D
% Cells annotated as TR3-56
% Cells in cluster
E
DiaT0 vs. Ctrl
DiaT1 vs. Ctrl
-Log10 p-val adj.
Avg. Log2 Fold Change
F
FlowAI Good Events
Lymphocytes
Single cells
Live
Panel 1
NK vs. CD3+
CD8+ vs. CD4+
Naive vs. AgExp
CD8low
Panel 2
G
Lymphocytes
Single cells
Live CD3+ cells
CD8+ vs. CD4+
Panel1
Panel2
H
CD8low
% from CD3+
0.16
NonDia
T1D

**Figure S12.**

Analysis of gene segment usage of TRAV (A), TRAJ (B), TRBV (C) and TRBJ (D) genes in conventional $CD8^+$ T cells from healthy (blue) and T1D (red) donors. TCR repertoires were profiled using 10x Immune Profiling with Feature Barcoding Technology. Dot represents mean, whiskers range from min to max.

Figure S12

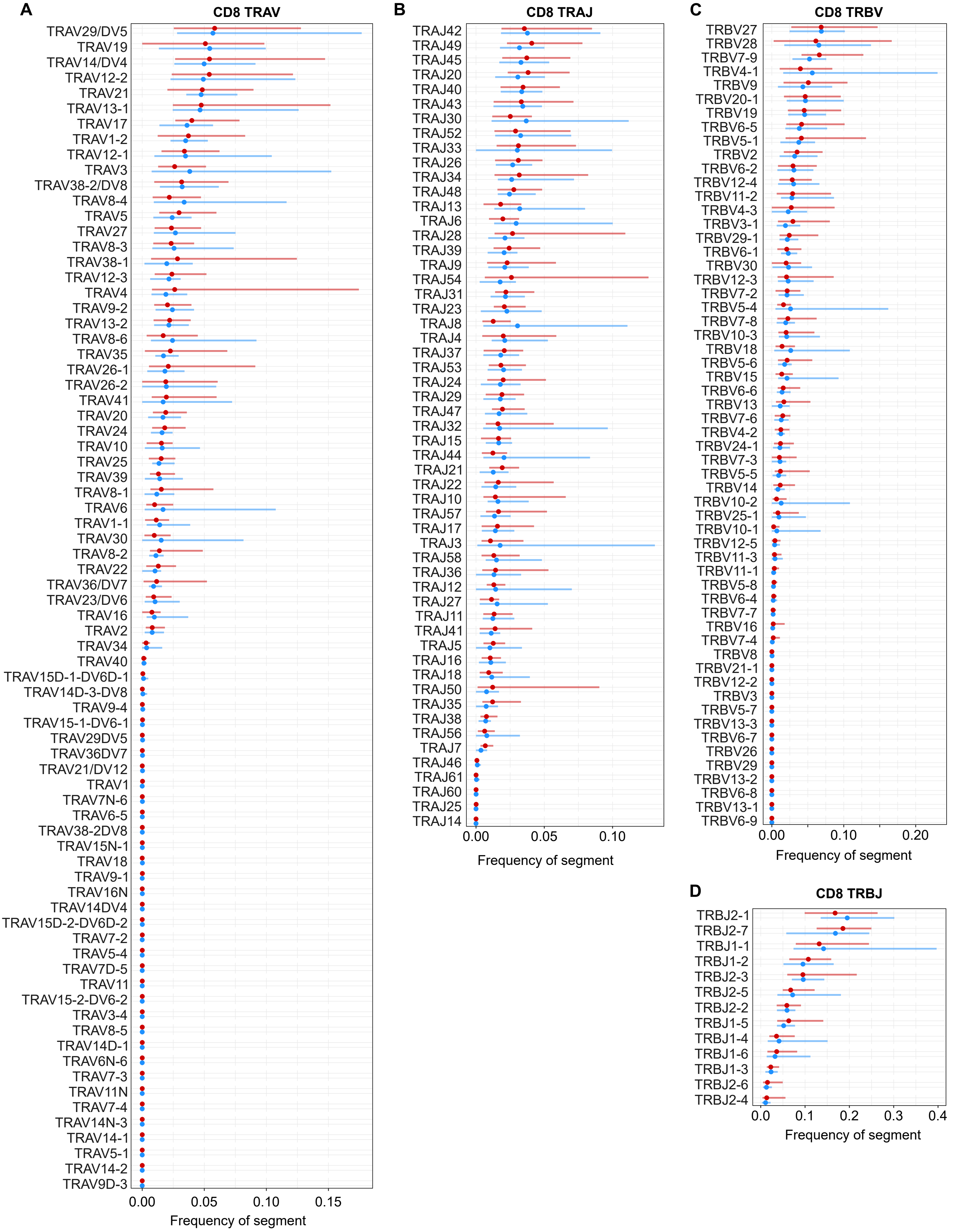

A
CD8 TRAV
TRAV29/DV5
TRAV19
TRAV14/DV4
TRAV12-2
TRAV21
TRAV13-1
TRAV17
TRAV1-2
TRAV12-1
TRAV3
TRAV38-2/DV8
TRAV8-4
TRAV5
TRAV27
TRAV8-3
TRAV38-1
TRAV12-3
TRAV4
TRAV9-2
TRAV13-2
TRAV8-6
TRAV35
TRAV26-1
TRAV26-2
TRAV41
TRAV20
TRAV24
TRAV10
TRAV25
TRAV39
TRAV8-1
TRAV6
TRAV1-1
TRAV30
TRAV8-2
TRAV22
TRAV36/DV7
TRAV23/DV6
TRAV16
TRAV2
TRAV34
TRAV40
TRAV15D-1-DV6D-1
TRAV14D-3-DV8
TRAV9-4
TRAV15-1-DV6-1
TRAV29DV5
TRAV36DV7
TRAV21/DV12
TRAV1
TRAV7N-6
TRAV6-5
TRAV38-2DV8
TRAV15N-1
TRAV18
TRAV9-1
TRAV16N
TRAV14DV4
TRAV15D-2-DV6D-2
TRAV7-2
TRAV5-4
TRAV7D-5
TRAV11
TRAV15-2-DV6-2
TRAV3-4
TRAV8-5
TRAV14D-1
TRAV6N-6
TRAV7-3
TRAV11N
TRAV7-4
TRAV14N-3
TRAV14-1
TRAV5-1
TRAV14-2
TRAV9D-3
0.00
0.05
0.10
0.15
Frequency of segment
B
CD8 TRAJ
TRAJ42
TRAJ49
TRAJ45
TRAJ20
TRAJ40
TRAJ43
TRAJ30
TRAJ52
TRAJ33
TRAJ26
TRAJ34
TRAJ48
TRAJ13
TRAJ6
TRAJ28
TRAJ39
TRAJ9
TRAJ54
TRAJ31
TRAJ23
TRAJ8
TRAJ4
TRAJ37
TRAJ53
TRAJ24
TRAJ29
TRAJ47
TRAJ32
TRAJ15
TRAJ44
TRAJ21
TRAJ22
TRAJ10
TRAJ57
TRAJ17
TRAJ3
TRAJ58
TRAJ36
TRAJ12
TRAJ27
TRAJ11
TRAJ41
TRAJ5
TRAJ16
TRAJ18
TRAJ50
TRAJ35
TRAJ38
TRAJ56
TRAJ7
TRAJ46
TRAJ61
TRAJ60
TRAJ25
TRAJ14
0.00
0.05
0.10
Frequency of segment
C
CD8 TRBV
TRBV27
TRBV28
TRBV7-9
TRBV4-1
TRBV9
TRBV20-1
TRBV19
TRBV6-5
TRBV5-1
TRBV2
TRBV6-2
TRBV12-4
TRBV11-2
TRBV4-3
TRBV3-1
TRBV29-1
TRBV6-1
TRBV30
TRBV12-3
TRBV7-2
TRBV5-4
TRBV7-8
TRBV10-3
TRBV18
TRBV5-6
TRBV15
TRBV6-6
TRBV13
TRBV7-6
TRBV4-2
TRBV24-1
TRBV7-3
TRBV5-5
TRBV14
TRBV10-2
TRBV25-1
TRBV10-1
TRBV12-5
TRBV11-3
TRBV11-1
TRBV5-8
TRBV6-4
TRBV7-7
TRBV16
TRBV7-4
TRBV8
TRBV21-1
TRBV12-2
TRBV3
TRBV5-7
TRBV13-3
TRBV6-7
TRBV26
TRBV29
TRBV13-2
TRBV6-8
TRBV13-1
TRBV6-9
0.00
0.10
0.20
Frequency of segment
D
CD8 TRBJ
TRBJ2-1
TRBJ2-7
TRBJ1-1
TRBJ1-2
TRBJ2-3
TRBJ2-5
TRBJ2-2
TRBJ1-5
TRBJ1-4
TRBJ1-6
TRBJ1-3
TRBJ2-6
TRBJ2-4
0.0
0.1
0.2
0.3
0.4
Frequency of segment

**Figure S13.**

Analysis of gene the segment usage of TRAV (A), TRAJ (B), TRBV (C) and TRBJ (D) genes in conventional $CD4^+$ T cells from healthy (blue) and T1D (red) donors. TCR repertoires were profiled using 10x Immune Profiling with Feature Barcoding Technology. Dot represents mean, whiskers range from min to max.

Figure S13

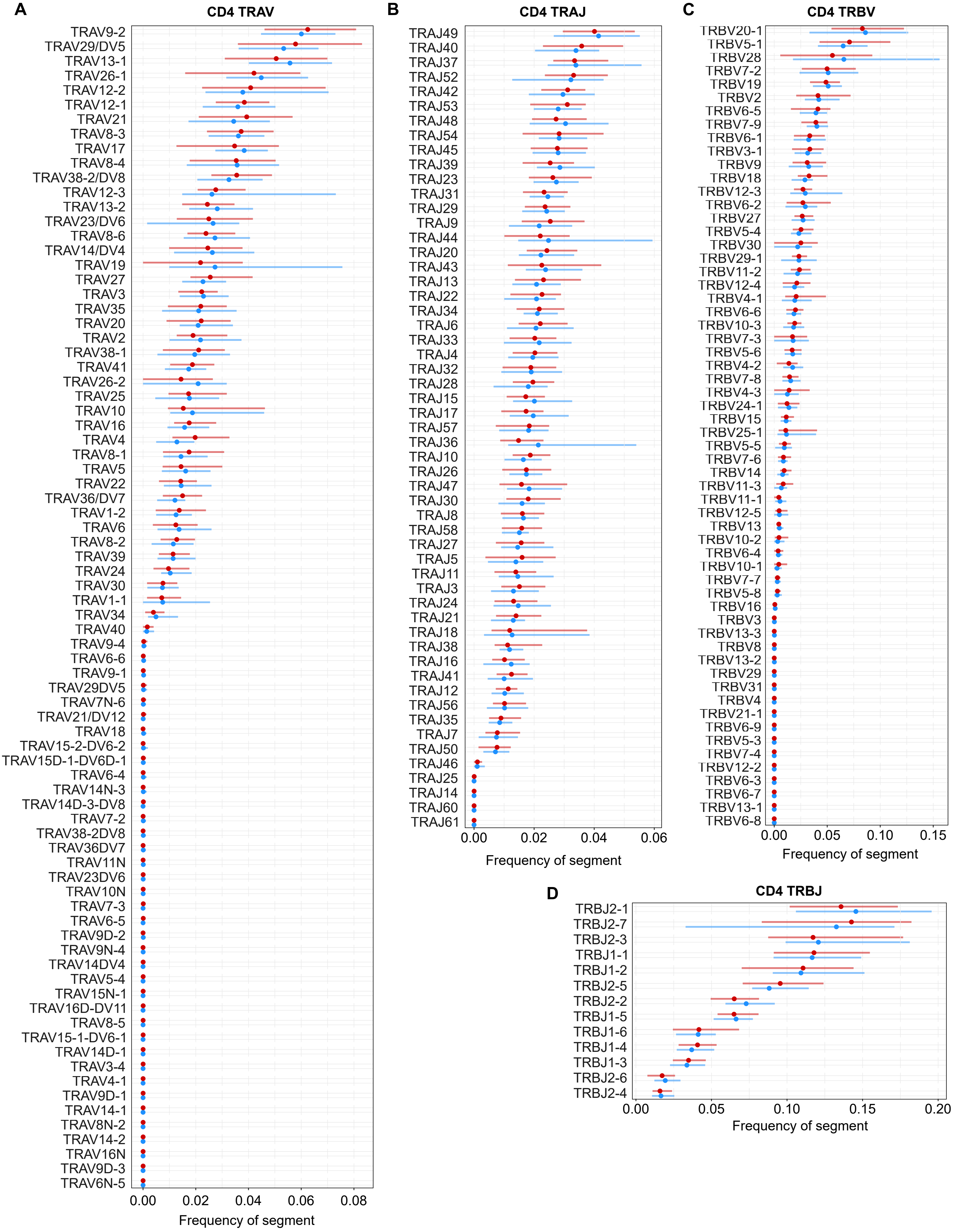
A
CD4 TRAV
TRAV9-2
TRAV29/DV5
TRAV13-1
TRAV26-1
TRAV12-2
TRAV12-1
TRAV21
TRAV8-3
TRAV17
TRAV8-4
TRAV38-2/DV8
TRAV12-3
TRAV13-2
TRAV23/DV6
TRAV8-6
TRAV14/DV4
TRAV19
TRAV27
TRAV3
TRAV35
TRAV20
TRAV2
TRAV38-1
TRAV41
TRAV26-2
TRAV25
TRAV10
TRAV16
TRAV4
TRAV8-1
TRAV5
TRAV22
TRAV36/DV7
TRAV1-2
TRAV6
TRAV8-2
TRAV39
TRAV24
TRAV30
TRAV1-1
TRAV34
TRAV40
TRAV9-4
TRAV6-6
TRAV9-1
TRAV29DV5
TRAV7N-6
TRAV21/DV12
TRAV18
TRAV15-2-DV6-2
TRAV15D-1-DV6D-1
TRAV6-4
TRAV14N-3
TRAV14D-3-DV8
TRAV7-2
TRAV38-2DV8
TRAV36DV7
TRAV11N
TRAV23DV6
TRAV10N
TRAV7-3
TRAV6-5
TRAV9D-2
TRAV9N-4
TRAV14DV4
TRAV5-4
TRAV15N-1
TRAV16D-DV11
TRAV8-5
TRAV15-1-DV6-1
TRAV14D-1
TRAV3-4
TRAV4-1
TRAV9D-1
TRAV14-1
TRAV8N-2
TRAV14-2
TRAV16N
TRAV9D-3
TRAV6N-5
0.00
0.02
0.04
0.06
0.08
Frequency of segment
B
CD4 TRAJ
TRAJ49
TRAJ40
TRAJ37
TRAJ52
TRAJ42
TRAJ53
TRAJ48
TRAJ54
TRAJ45
TRAJ39
TRAJ23
TRAJ31
TRAJ29
TRAJ9
TRAJ44
TRAJ20
TRAJ43
TRAJ13
TRAJ22
TRAJ34
TRAJ6
TRAJ33
TRAJ4
TRAJ32
TRAJ28
TRAJ15
TRAJ17
TRAJ57
TRAJ36
TRAJ10
TRAJ26
TRAJ47
TRAJ30
TRAJ8
TRAJ58
TRAJ27
TRAJ5
TRAJ11
TRAJ3
TRAJ24
TRAJ21
TRAJ18
TRAJ38
TRAJ16
TRAJ41
TRAJ12
TRAJ56
TRAJ35
TRAJ7
TRAJ50
TRAJ46
TRAJ25
TRAJ14
TRAJ60
TRAJ61
0.00
0.02
0.04
0.06
Frequency of segment
C
CD4 TRBV
TRBV20-1
TRBV5-1
TRBV28
TRBV7-2
TRBV19
TRBV2
TRBV6-5
TRBV7-9
TRBV6-1
TRBV3-1
TRBV9
TRBV18
TRBV12-3
TRBV6-2
TRBV27
TRBV5-4
TRBV30
TRBV29-1
TRBV11-2
TRBV12-4
TRBV4-1
TRBV6-6
TRBV10-3
TRBV7-3
TRBV5-6
TRBV4-2
TRBV7-8
TRBV4-3
TRBV24-1
TRBV15
TRBV25-1
TRBV5-5
TRBV7-6
TRBV14
TRBV11-3
TRBV11-1
TRBV12-5
TRBV13
TRBV10-2
TRBV6-4
TRBV10-1
TRBV7-7
TRBV5-8
TRBV16
TRBV3
TRBV13-3
TRBV8
TRBV13-2
TRBV29
TRBV31
TRBV4
TRBV21-1
TRBV6-9
TRBV5-3
TRBV7-4
TRBV12-2
TRBV6-3
TRBV6-7
TRBV13-1
TRBV6-8
0.00
0.05
0.10
0.15
Frequency of segment
D
CD4 TRBJ
TRBJ2-1
TRBJ2-7
TRBJ2-3
TRBJ1-1
TRBJ1-2
TRBJ2-5
TRBJ2-2
TRBJ1-5
TRBJ1-6
TRBJ1-4
TRBJ1-3
TRBJ2-6
TRBJ2-4
0.00
0.05
0.10
0.15
0.20
Frequency of segment
